# Direct photoresponsive inhibition of a p53-like transcription activation domain in PIF3 by *Arabidopsis* phytochrome B

**DOI:** 10.1101/2021.05.11.443493

**Authors:** Chan Yul Yoo, Qing Sang, Jiangman He, Yongjian Qiu, Lingyun Long, Ruth Jean-Ae Kim, Emily G. Chong, Joseph Hahm, Nicholas Morffy, Pei Zhou, Lucia C. Strader, Akira Nagatani, Beixin Mo, Xuemei Chen, Meng Chen

## Abstract

Phytochrome B (PHYB) triggers diverse light responses in *Arabidopsis* by binding to a group of antagonistically acting PHYTOCHROME-INTERACTING transcription FACTORs (PIFs) to promote PIF degradation, consequently downregulating PIF target genes. However, whether PHYB directly controls the transactivation activity of PIFs remains ambiguous. Here we show that the prototypic PIF, PIF3, possesses a p53-like transcription activation domain (TAD) consisting of a sequence-specific, hydrophobic activator motif surrounded by acidic residues. A PIF3mTAD mutant in which the activator motif is replaced with alanines fails to activate PIF3 target genes in *Arabidopsis* in dark, light, and shade conditions, validating the *in vivo* functions of the PIF3 TAD. Intriguingly, binding of PHYB’s N-terminal photosensory module to the PHYB-binding site adjacent to the TAD inhibits its transactivation activity. These results unveil a photoresponsive transcriptional switching mechanism in which photoactivated PHYB directly masks the transactivation activity of PIF3. Our study also suggests the unexpected conservation of sequence-specific TADs between the animal and plant kingdoms.

## INTRODUCTION

The plant genome is exquisitely sensitive to and can be transcriptionally reprogrammed by changes in light intensity and composition^1, 2^. In particular, alterations in the ratio of red (R, 660 nm) and far-red (FR, 730 nm) light are powerful environmental cues that inform about space and time – such as the availability of photosynthetically active R-light radiation; the timing of sunrise and sunset, as twilight is enriched in FR light; and the threat by neighboring vegetation or shade, which depletes R light^2, 3^. Plants detect R and FR light through evolutionarily conserved photoreceptors called phytochromes (PHYs)^4^. The prototypical plant PHY is a homodimer: each monomer comprises an N-terminal photosensory module and a C-terminal signaling output module^5, 6^. The N-terminal photosensory module consists of four domains: an N-terminal extension, a PAS (Period-Arnt-Single-Minded) domain, a GAF (cGMP phosphodiesterase/adenylate cyclase/FhlA) domain that binds a tetrapyrrole chromophore, and a PHY (phytochrome-specific) domain. At the PAS-GAF interface, there is an unusual figure-eight knot called the “light-sensing knot”, which is tightly linked to the chromophore and plays an important role in signaling^7–9^. The C-terminal output module contains two tandem PAS domains and a histidine kinase-related domain (HKRD), which collectively mediate dimerization, subcellular localization, and signaling^10–13^. Absorption of R or FR light by the tetrapyrrole moiety embedded in the N-terminal photosensory module photoconverts PHYs between two relatively stable forms: the R-light-absorbing inactive Pr and the FR-light-absorbing active Pfr^5, 6^. The conformational changes of PHYs dictate their subcellular localization: while the Pr localizes in the cytoplasm, the Pfr accumulates in the nucleus and forms PHY-containing subnuclear foci called photobodies, concomitantly reprogramming the expression of hundreds of light-responsive genes^14, 15^. As such, the equilibrium of the Pr and Pfr of PHYs quantitatively connects environmental light cues to light-responsive gene expression. PHY-mediated gene regulation is at the center of plant-environment interactions and profoundly impacts all aspects of plant development, growth, metabolism, and immunity, enabling plants to thrive in complex natural environments^16^.

In *Arabidopsis*, PHYs are encoded by a small gene family, *PHYA*-*E*^17^. PHYB is the most prominent PHY, evidently because only PHYB directly binds to an entire family of basic/helix-loop-helix (bHLH) transcription factors named PHYTOCHROME-INTERACTING FACTORs (PIFs)^18^. The PIF family comprises eight members: PIF1-8 (PIF2 and PIF6 are also called PIL1 and PIL2 [PIF3-Like1 and 2], respectively)^19, 20^. The C-termini of all PIFs contain a bHLH domain for dimerization and DNA binding to the G-box as well as a PIF-binding E-box (PBE-box)^21, 22^, while the N-termini contain an Active-PHYB Binding (APB) motif, which interacts preferentially with photoactivated PHYB, specifically the light-sensing knot in the N-terminal photosensory module^9, 18^. PIF1 and PIF3 also interact with the Pfr form of PHYA via a separate Active-PHYA Binding (APA) motif, which is absent in the rest of the PIFs^20^. In general, PIFs play antagonistic roles in PHYB signaling^23, 24^. During seedling development, after seed germination underground in the absence of light, four PIFs, PIF1, PIF3, PIF4, and PIF5, promote the dark-grown developmental program called skotomorphogenesis by blocking leaf development and chloroplast biogenesis and instead accelerating the elongation of the embryonic stem (hypocotyl), a coordinated body plan allowing seedlings to quickly emerge from the soil^25, 26^. To promote hypocotyl elongation, PIFs bind directly to the enhancer regions of growth-relevant genes, such as those involved in the biosynthesis and signaling of the plant growth hormone auxin, and activate their transcription^21, 22^. Upon seedling emergence from the ground and exposure to light, photoactivated PHYB accumulates in the nucleus and interacts with PIF1, PIF3, PIF4, and PIF5 to promote their rapid phosphorylation, ubiquitylation, and degradation by the proteasome, thereby reprogramming the expression of light-responsive PIF target genes^13, 27–29^. PHYB-triggered degradation of PIFs has been considered a central mechanism of PHYB signaling^20^, implying that PHYB regulates the transcription of PIF target genes mainly indirectly by controlling the abundance of PIFs.

Accumulating circumstantial evidence suggests that PHYB may also directly control the transcriptional activity of PIFs. First, not all PIFs are degraded in the light or by photoactivated PHYB. For example, the protein level of PIF7 is not significantly different between dark and light conditions^30^. Also, although PIF4 and PIF5 rapidly diminish during initial exposure to light, when seedlings are grown under diurnal or continuous light conditions, both can accumulate to high levels during the daytime, when PHYB is active^31, 32^. Moreover, the stability of PIFs varies at different temperatures, despite being almost undetectable in ambient temperatures, PIF3 becomes stabilized in the light in cold temperatures^33^. Therefore, the PHYB-PIF interaction could regulate the activity of the photostable PIFs. Second, although PHYB triggers the light-induced degradation of PIF3, recent studies surprisingly revealed that the light-dependent interaction between the light-sensing knot of PHYB and the APB of PIF3 is not responsible for PIF3 degradation^10, 34, 35^. Instead, PIF3 degradation relies on a weaker, light-independent interaction with PHYB’s C-terminal output module^10, 34, 35^. These results leave open the important question of what is the functional significance of the PHYB-APB interaction in PIF3 regulation. One proposed hypothesis is that binding of PHYB to the APB regulates the transcriptional activity of PIFs^34, 36^. Supporting this hypothesis, the PHYB-APB interaction was shown to attenuate PIF3’s DNA-binding activity independently of its degradation^34, 36^. However, because the APB motif is located at the very N-terminus of PIF3, far from the C-terminal bHLH domain and PHYB does not interact directly with the bHLH^18, 21^, it remains unclear whether PHYB influences the DNA binding of PIF3 directly or indirectly via the control of its transactivation activity.

The main roadblock that hinders our understanding of the relationship between PHYB and the transactivation activity of PIFs is the lack of a molecular understanding of PIFs’ transcription activation domains (TADs). It is well understood that transcriptional activators follow a modular design consisting of a DNA-binding domain to selectively bind to specific DNA sequences (enhancers) and a TAD to initiate transcription^37^. Although the classes of DNA-binding domains have been well defined and can be recognized based on conserved amino acid sequences^38^, TADs remain difficult to predict due to their poor sequence conservation^39–41^. As a result, despite PIFs being widely accepted as transcriptional activators^22^, their TAD sequences have not been precisely defined. A previous attempt to map the TAD of PIF3 identified two separate regions in its N-terminal half that carry transactivation activity in yeast, although these putative TADs have not been validated *in planta*^42^. To further explore the possibility of a direct link between PHYB and the transactivation activity of PIFs, first, we utilized truncation analysis and alanine-scanning mutagenesis combined with yeast and *in planta* assays to determine the TAD of PIF3. Our results show that PIF3 possesses a single TAD, which surprisingly resembles the TADs of the mammalian tumor suppressor p53 and the yeast activator Gcn4, revealing the unexpected conservation of sequence-specific TADs across the animal, fungal and plant kingdoms. The PIF3 TAD is conserved to various degrees among PIF3 paralogs. Furthermore, we demonstrate that binding of PHYB’s N-terminal photosensory module to the APB motif of PIF3 represses PIF3’s transactivation activity, indicating a direct role for PHYB in the photoinhibition of the transactivation activity of PIFs.

## RESULTS

### The aa_91-114_ region confers PIF3’s TAD activity in yeast

Ideally, defining and validating the TAD of PIF3 requires evaluating the *in planta* functions of a PIF3 mutant specifically lacking its TAD activity while retaining its other biochemical functions. A previous study used a modified yeast one-hybrid screen and random mutagenesis to identify two regions of PIF3 with transactivation activity in yeast^42^. However, the PIF3 mutants isolated via this approach contained multiple mutations; therefore, the residues directly and specifically required for the TAD activity could not be unambiguously pinpointed^42^. To circumvent these potential issues, we chose to systematically map the region(s) and residues that are both necessary and sufficient for PIF3’s transactivation activity by truncation analysis and mutagenesis. Because PIF3 has been shown to be a potent activator in yeast^42^, we first used yeast transactivation assays to examine PIF3’s TAD activity; we fused the Gal4 DNA-binding domain (DBD) to either the full-length PIF3 or a series of N- or C-terminal truncation fragments in the bait construct of the Matchmaker Gold yeast two-hybrid system (Fig. 1a). We evaluated the self-activation of the recombinant DBD-PIF3 proteins using two assays: (1) a cell viability assay testing the activation of the Aureobasidin A (AbA) resistance gene *AUR1-C* and (2) a liquid assay examining the activation of a second reporter, β-galactosidase (Fig. 1a). Interestingly, all PIF3 fragments containing the region between amino acids 91 and 114 (aa_91-114_), i.e., PIF3-N1 to PIF3-N4, as well as PIF3-C1 and PIF3-C2, retained at least 87% of the activity of full-length PIF3 (Fig. 1a). In contrast, the two fragments missing part of aa_91-114_, i.e., PIF3-N5 and PIF3-C3, completely lacked the activity (Fig. 1a). The aa_91-114_ fragment alone (PIF3-M2) was sufficient to activate gene expression, albeit with reduced activity (Fig. 1a). The weaker activity of aa_91-114_ could be due to a lack of structural support provided by the surrounding protein context because a larger fragment containing amino acids 76 to 114 (PIF3-M1) displayed significantly higher activity (Fig. 1a).

**Fig. 1.**
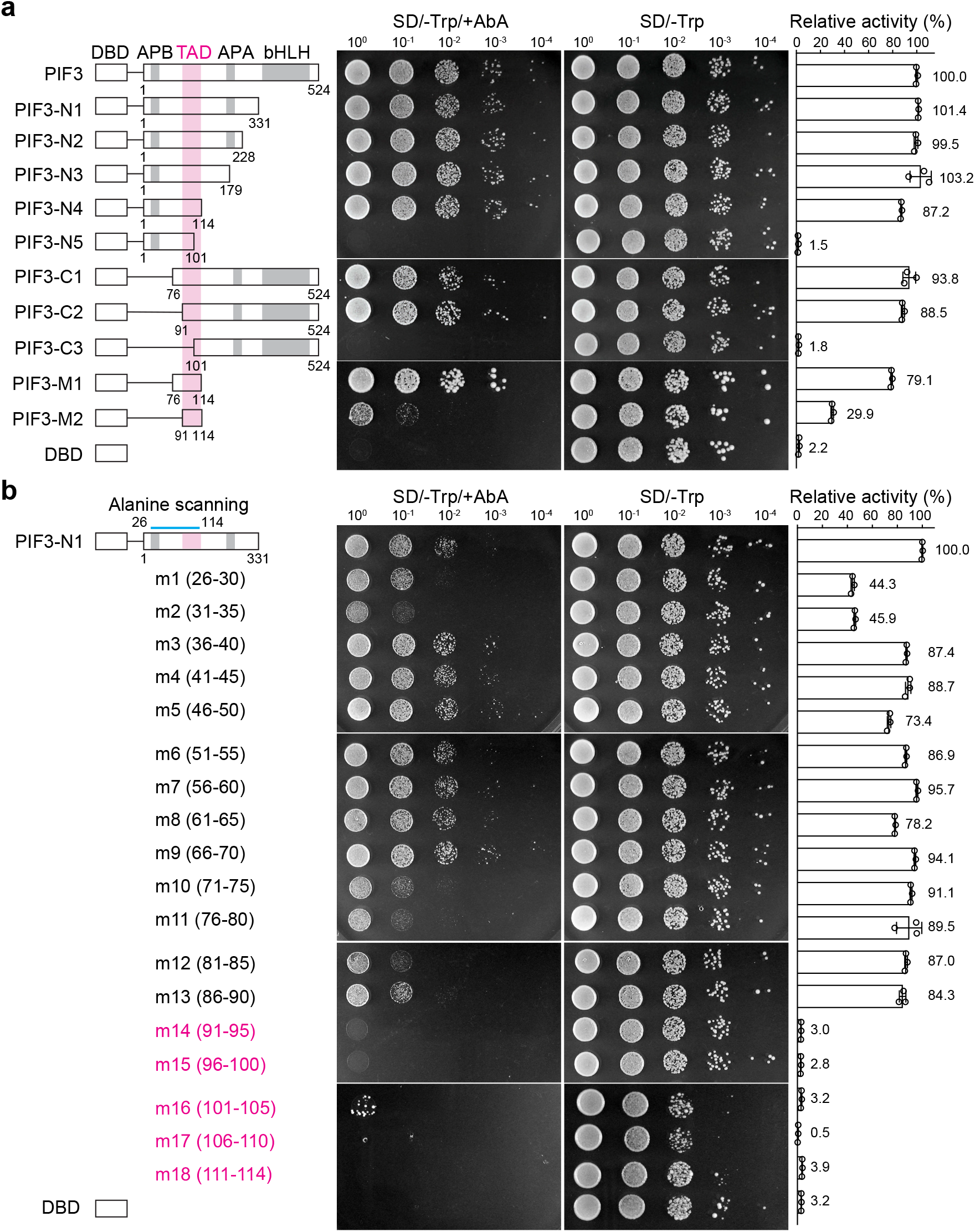
The aa_91-114_ region confers the TAD activity of PIF3 in yeast. **a** PIF3 and a series of PIF3 truncation fragments were fused to the Gal4 DNA binding domain (DBD), as shown in the schematics, using the bait vector of the yeast two-hybrid system and evaluated for their transactivation activity. The magenta column highlights the aa_91-114_ region. APB, active PHYB binding region; APA, active PHYA binding region. **b** Identification of the residues necessary for PIF3’s TAD activity via alanine scanning mutagenesis. A series of alanine-scanning mutants (m1-m18) were generated between amino acids 26 and 114 (indicated by the blue bar) in the PIF3-N1 construct. The range of the mutated amino acids for each mutant is shown in parentheses. The mutants that lost their transactivation activity are highlighted in magenta. **a**, **b** The middle panel shows serial dilutions of the yeast strains containing the respective constructs grown on either SD/-Trp/+AbA or SD/-Trp (control) media. The right panel shows the relative transactivation activities quantified using the yeast liquid β-galactosidase assay. The transactivation activities were calculated relative to that of either PIF3 **(a)** or PIF3-N1 **(b)**. DBD alone was used as a negative control. Error bars represent the s.d. of three biological replicates; the numbers represent the mean value of the relative activity.

Our results of aa_91-114_ being an activating region is consistent with the previous study, which showed that the slightly larger aa_90-120_ region carries transactivation activity^42^. However, we did not detect any transactivation activity between amino acids 27 and 43, which encompass the second TAD suggested by the previous study^42^. None of the four N-terminal fragments of PIF3, aa_1-52_, aa_1-76_, aa_1-90_, and aa_1-101_, showed any transactivation activity in our assays (Fig. 1a and Supplementary Fig. 1). To further resolve this discrepancy, we performed alanine-scanning mutagenesis in the region between amino acids 26 and 114 in the PIF3-N1 (aa_1-331_) fragment by substituting every 5 consecutive amino acids with alanines (Fig. 1b). Corroborating the results of the truncation analysis, only the alanine substitution mutants in the aa_91-114_ region (m14 to m18) completely abrogated the transactivation activity (Fig. 1b). In contrast, all mutants carrying mutations within the suggested second activating region (m1 to m4) remained active, despite m1 and m2 showing reduced activities (Fig. 1b), indicating that the aa_26-43_ region is not essential for PIF3’s transactivation activity. Together, the results of the truncation analysis and alanine-scanning mutagenesis indicate that aa_91-114_ is the only region both required and sufficient for PIF3’s transactivation activity and therefore confers PIF3’s TAD function in yeast.

### PIF3 possesses a p53-like TAD

Studies of animal and yeast activators suggest that one class of TADs comprises a short, sequence-specific activator motif of hydrophobic residues embedded in an intrinsically disordered region enriched in acidic residues^40, 41, 43–45^. One identified activator motif is ΦxxΦΦ, where Φ indicates a bulky hydrophobic residue and x is any other amino acid^40^. For example, both TADs in the mammalian tumor suppressor p53, AD1 and AD2, contain the ΦxxΦΦ motif^43^. The same motif is also found in the yeast activator Gcn4^40, 41^. Surprisingly, aligning the PIF3 TAD to the TADs of p53 and Gcn4 revealed striking similarities: PIF3’s TAD features a **F**VP**WL** motif flanked by aspartate and glutamate residues (Fig. 2a). The FVPWL motif in PIF3 shares the same hydrophobic residues as the activator motifs in p53 and Gcn4, including the two bulky aromatic amino acids F93 and W96. Both the activator motif and the negatively charged residues have been shown to play important roles in transactivation. For example, the ΦxxΦΦ motif in p53 AD1 and AD2 mediates hydrophobic interactions with the transcriptional coactivator cyclic-AMP response element-binding protein (CREB)-binding protein (CBP) and its paralog p300^43^. The surrounding acidic residues participate in electrostatic interactions with transcriptional coactivators and create an intrinsically disordered context in the unbound state to enhance the accessibility of the activator motif^44^. Both the ΦxxΦΦ motif and surrounding acidic residues are highly conserved in PIF3 orthologs in eudicots (Fig. 2a), supporting the idea that PIF3 possesses a p53-like TAD.

**Fig. 2.**
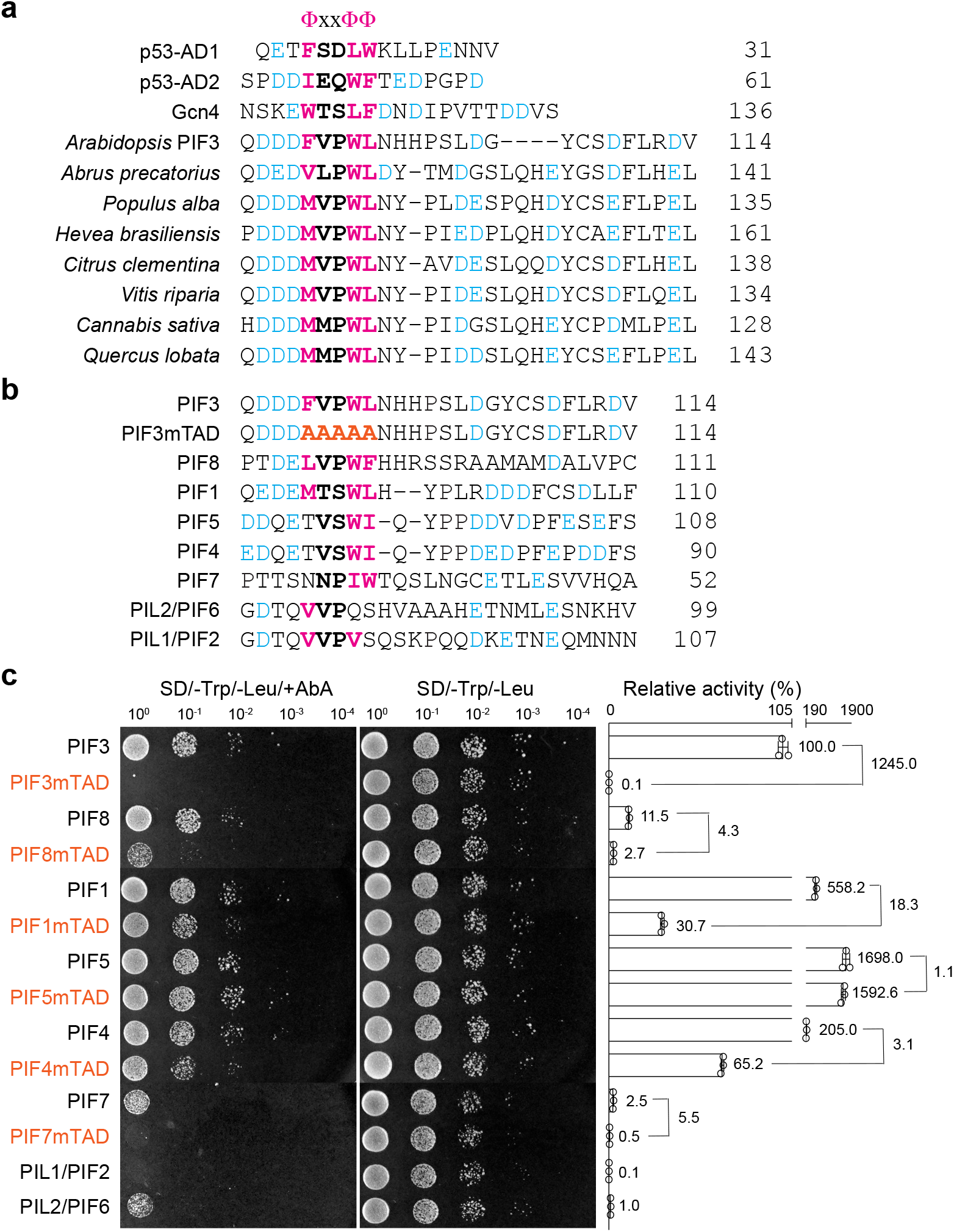
PIF3 possesses a p53-like TAD. **a** Amino acid sequence alignment of the TADs of *Homo sapiens* p53 (BAC16799.1), *Saccharomyces cerevisiae* Gcn4 (QHB08060.1), and select PIF3 orthologs from eudicots, including *Arabidopsis thaliana* (NP_001318964.1), *Abrus precatorius* (XP_027363600.1), *Populus alba* (XP_034896271.1), *Hevea brasiliensis* (XP_021639209.1), *Citrus clementina* (XP_006423962.1), *Vitis riparia* (XP_034707239.1), *Cannabis sativa* (XP_030504594.1), and *Quercus lobata* (XP_030957900.1). **b** Amino acid sequence alignment of the TAD regions of PIF3, PIF3mTAD, and the other PIF paralogs in *Arabidopsis*. The substituted alanines in PIF3mTAD are labeled in orange. **a, b** The conserved activator ΦxxΦΦ motifs are highlighted in bold; the critical hydrophobic residues in the ΦxxΦΦ motif and surrounding acidic residues are labeled in magenta and blue, respectively. **c** Yeast transactivation assays showing the transactivation activities of the full-length PIF3 and PIF3 paralogs in *Arabidopsis* and the respective mTAD mutants of PIF3, PIF8, PIF1, PIF5, PIF4, and PIF7. All mTAD mutants contain five alanine substitutions in the respective ΦxxΦΦ motifs. The left panel shows serial dilutions of the yeast strains containing the respective constructs grown on either SD/-Trp/-Leu/+AbA or SD/-Trp/-Leu (control) media. The right panel shows the relative transactivation activities quantified using the yeast liquid β-galactosidase assay. The yeast assays were repeated five times; the results of a representative experiment are shown. The transactivation activities were calculated relative to that of PIF3. Error bars represent the s.d. of three technical replicates; the numbers represent the mean value of the relative activity. Fold changes in the activity of mTAD mutants relative to their respective wild-type proteins are denoted.

To evaluate the contribution of the conserved residues around the ΦxxΦΦ motif to PIF3 transactivation activity, we mutated amino acids 91 to 100 individually to alanines in the PIF3-N1 fragment. The single-amino-acid substitution mutants, however, had little effect on PIF3’s TAD activity (Supplementary Fig. 2a, b), suggesting that the main functional and/or structural attributes of the TAD are fulfilled redundantly by multiple residues. We then replaced the five residues of the FVPWL motif with alanines (a combination that was not included in the alanine-scanning mutagenesis) in full-length PIF3 and named it PIF3mTAD (Fig. 2b). PIF3mTAD completely lost its transactivation activity, confirming that this is the only TAD in PIF3 (Fig. 2c). In addition, replacing only the three key hydrophobic residues in the ΦxxΦΦ motif, namely F93, W96, and L97, with arginine or serine also abolished TAD activity (Supplementary Fig. 2a, c); similar mutations were shown to impede the activity of p53^46^, suggesting that the critical roles of these hydrophobic residues in TAD activity are conserved in PIF3.

The TAD region of PIF3 was previously recognized as MUF1 (motif of unknown function 1), which is moderately conserved among PIF family members (Fig. 2b)^47^. The ΦxxΦΦ motif can be recognized in PIF1, PIF4, PIF5, PIF7, and PIF8. While PIF1 and PIF8 contain a perfectly composed ΦxxΦΦ motif, PIF4, PIF5, and PIF7 are missing the initial hydrophobic residue (Fig. 2b). The surrounding acidic residues are conserved to various degrees among PIF1, PIF4, PIF5, PIF7, and PIF8. In PIF1, PIF4, and PIF5, multiple acidic residues are present on both sides of the ΦxxΦΦ motif, whereas, in PIF7 and PIF8, at least one side of the ΦxxΦΦ motif contains significantly fewer acidic residues (Fig. 2b). The attributes of the PIF3 TAD are least conserved in PIL1/PIF2 and PIL2/PIF6; the ΦxxΦΦ motif is barely recognizable and missing the potentially critical aromatic amino acids (Fig. 2b). We then examined the transactivation activities of all PIF members in yeast by fusing Gal4 DBD to the full-length PIF sequences. The PIF3 paralogs displayed a wide range of transactivation activities (Fig. 2c). The differences in activity could be, at least in part, explained by the degree of conservation of the two key attributes of the TAD. For example, PIF1, PIF4, PIF5, PIF7, and PIF8, which contain recognizable attributes of the PIF3 TAD, exhibited clear transactivation activity (Fig. 2b, c). PIF1, PIF4, and PIF5 were the most potent activators, with significantly higher transactivation activity than PIF3, whereas PIF7 and PIF8 were less active than PIF3 (Fig. 2c). Conversely, PIL1 and PIL2, which show little similarity to the PIF3 TAD in the corresponding region, had the least transactivation activity; PIL1 in particular showed no detectable activity (Fig. 2b, c). Interestingly, PIL1 has been suggested to heterodimerize with PIF4, PIF5, and PIF7 to negatively regulate their activities^48^. Therefore, the activities of the PIF family members might reflect their functional roles in PHY signaling.

To examine whether the ΦxxΦΦ motif contributes to the transactivation activities of PIF3 paralogs, we replaced the corresponding ΦxxΦΦ residues in PIF1, PIF4, PIF5, PIF7, and PIF8 with alanines to create their respective mTAD mutants. Intriguingly, only PIF7mTAD lost its transactivation activity, suggesting the ΦxxΦΦ region in PIF7 is its sole TAD (Fig. 2c). In contrast, mutating the ΦxxΦΦ motif did not abolish the activity of PIF1, PIF4, PIF5, or PIF8. PIF1mTAD, PIF4mTAD, and PIF8mTAD remained active but had dramatically reduced activities compared with their respective wild-type proteins; the activity of PIF5 was barely affected by the mTAD mutations (Fig. 2c). These results suggest that either the ΦxxΦΦ sequences in PIF1, PIF4, PIF5, and PIF8 are not essential for their TAD activity or these PIFs contain one or more TADs in addition to the one corresponding to the PIF3 TAD. Together, the current data indicate that neither the sequence nor the activity of the PIF3 TAD is perfectly conserved among PIF family members, implying a possible divergence in the mechanisms of transactivation and regulation among different PIFs. These differences warrant further in-depth investigations on the TAD functions of each individual PIF; we therefore focus the rest of the study on PIF3.

### Mutating the TAD does not affect PHYA or PHYB binding

The TAD of PIF3 lies between the APB and APA motifs (Fig. 1a)^18^. Next, we used *in vitro* GST pulldown assays to test whether the alanine mutations in PIF3mTAD had an effect on PIF3 binding to PHYB and PHYA. Similar to GST-PIF3, GST-PIF3mTAD preferentially bound to the active Pfr forms of hemagglutinin (HA)-tagged PHYB and PHYA without a detectable change in the pulldown results (Fig. 3). These results indicate that the ΦxxΦΦ activator motif of the PIF3 TAD does not directly engage in the interactions with PHYA and PHYB.

**Fig. 3.**
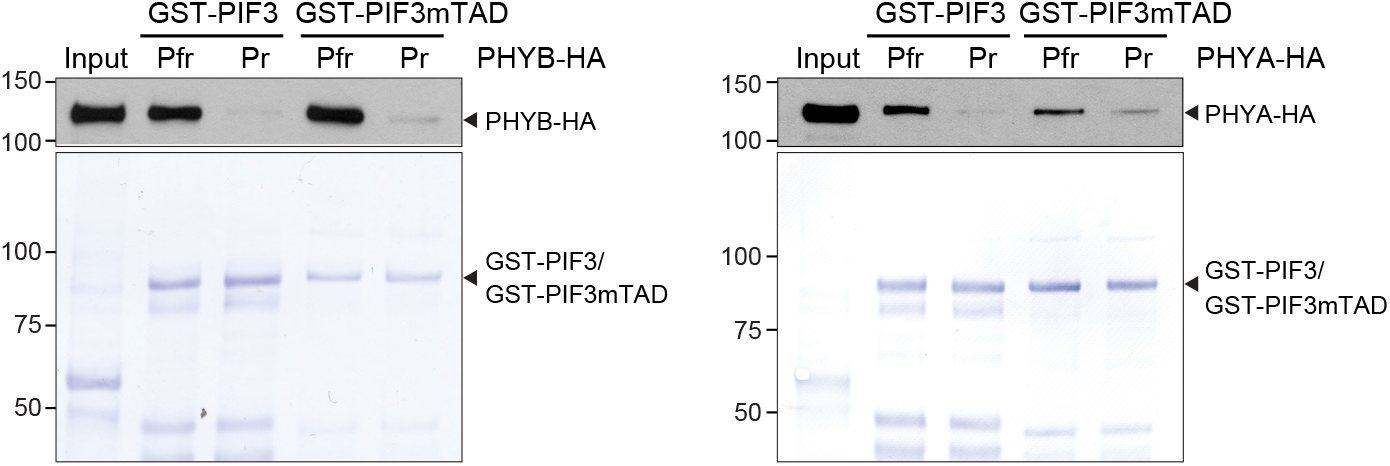
PIF3mTAD retains light-dependent interactions with PHYA and PHYB. GST pulldown results showing that PIF3mTAD, like PIF3, interacts preferentially with active PHYA and PHYB. GST pulldown assays were performed using recombinant GST-PIF3, GST-PIF3mTAD, or GST to pull down *in vitro* translated HA-tagged PHYA (PHYA-HA) or PHYB (PHYB-HA) conjugated with a phycocyanobilin chromophore. The pulldown assays were performed in either R or FR light to maintain PHYA-HA and PHYB-HA in the Pfr or Pr conformer, respectively. Input and bound PHYA-HA and PHYB-HA fractions were detected via immunoblots (upper panels) using anti-HA antibodies. Immobilized GST, GST-PIF3, and GST-PIF3mTAD are shown in the Coomassie Brilliant Blue-stained SDS-PAGE gels (lower panels).

### PIF3mTAD compromises PIF3’s functions in skotomorphogenesis

To validate the PIF3 TAD *in vivo*, we generated transgenic lines expressing HA- and YFP-tagged PIF3 (HA-YFP-PIF3) or PIF3mTAD (HA-YFP-PIF3mTAD) under the native *PIF3* promoter as described previously^49^. The transactivation activity of PIF3 was best demonstrated during skotomorphogenesis. In dark-grown *Arabidopsis* seedlings, PIF3 binds to the enhancer regions of its target genes to activate their expression and consequently promote skotomorphogenesis, such as exaggerated elongation of the hypocotyl^22, 50^. If the identified PIF3 TAD is the sole functional TAD in PIF3 *in vivo*, PIF3mTAD should be defective in PIF3 functions in skotomorphogenesis. However, skotomorphogenesis is maintained reduntantly by three additional PIF members, PIF1, PIF4, and PIF5, which hetero-dimerize or hetero-oligomerize with PIF3^15, 22, 24, 51^. Therefore, the potential effects of PIF3mTAD could be obscured in the presence of the other three PIFs. To circumvent this redundancy, we generated transgenic lines expressing HA-YFP-PIF3 or HA-YFP-PIF3mTAD in a *pif1pif3pif4pif5* quadruple mutant (*pifq*)^23^. For each construct, we picked two transgenic lines expressing similar levels of HA-YFP-PIF3 or HA-YFP-PIF3mTAD. Despite being controlled by the native *PIF3* promoter, the tagged PIF3 proteins in all four transgenic lines were expressed at higher levels than was endogenous PIF3 in the wild-type Col-0 (Fig. 4a). We therefore assessed the activity of HA-YFP-PIF3mTAD via comparison with that of HA-YFP-PIF3. Dark-grown *pifq* seedlings display de-etiolated phenotypes with short hypocotyls compared with Col-0 (Fig. 4b, c)^23, 24^. The two transgenic lines expressing HA-YFP-PIF3 (*PIF3/pifq 1-2* and *9-5*) fully rescued the short-hypocotyl phenotype of *pifq*. In contrast, the two lines expressing HA-YFP-PIF3mTAD (*PIF3mTAD/pifq 2-1* and *4-5*) only partially reverted the hypocotyl phenotype of *pifq* and remained significantly shorter than the Col-0 and the *PIF3/pifq* seedlings (Fig. 4b, c). Because HA-YFP-PIF3 and HA-YFP-PIF3mTAD were expressed at similar levels in the transgenic lines (Fig. 4a) and both localized to the nucleus (Fig. 4d), the discrepancy between the *PIF3/pifq* and *PIF3mTAD/pifq* lines is most likely due to the defect in the transactivation activity of HA-YFP-PIF3mTAD. Indeed, the steady-state transcript levels of four PIF3 target genes^22^, *PIL1*, *ATHB-2*, *XTR7* and *RD20*, were significantly lower in the *PIF3mTAD/pifq* lines than in the *PIF3/pifq* lines (Fig. 4e). To exclude the less-likely possibility that the reduced expression of the PIF3 target genes in the *PIF3mTAD/pifq* lines was due to a defect of PIF3mTAD in DNA binding, we examined the binding of HA-YFP-PIF3 and HA-YFP-PIF3mTAD to the G-boxes in the enhancer region of the marker gene *PIL1* via chromatin immunoprecipitation (ChIP). The results of the ChIP experiments confirmed that the mutations in PIF3mTAD do not affect its DNA binding activity (Fig. 4f). Together, these results support the conclusion that PIF3mTAD is defective in the transactivation activity of PIF3, providing strong *in vivo* evidence validating the aa_91-114_ region as PIF3’s TAD. The fact that the *PIF3mTAD/pifq* lines could still partially reverse the hypocotyl phenotype of *pifq* (Fig. 4a, b) suggests that either the PIF3mTAD mutations do not completely abolish the TAD activity *in vivo*, or alternatively, PIF3 can exert its activator functions through other associated transcription activators^49^.

**Fig. 4.**
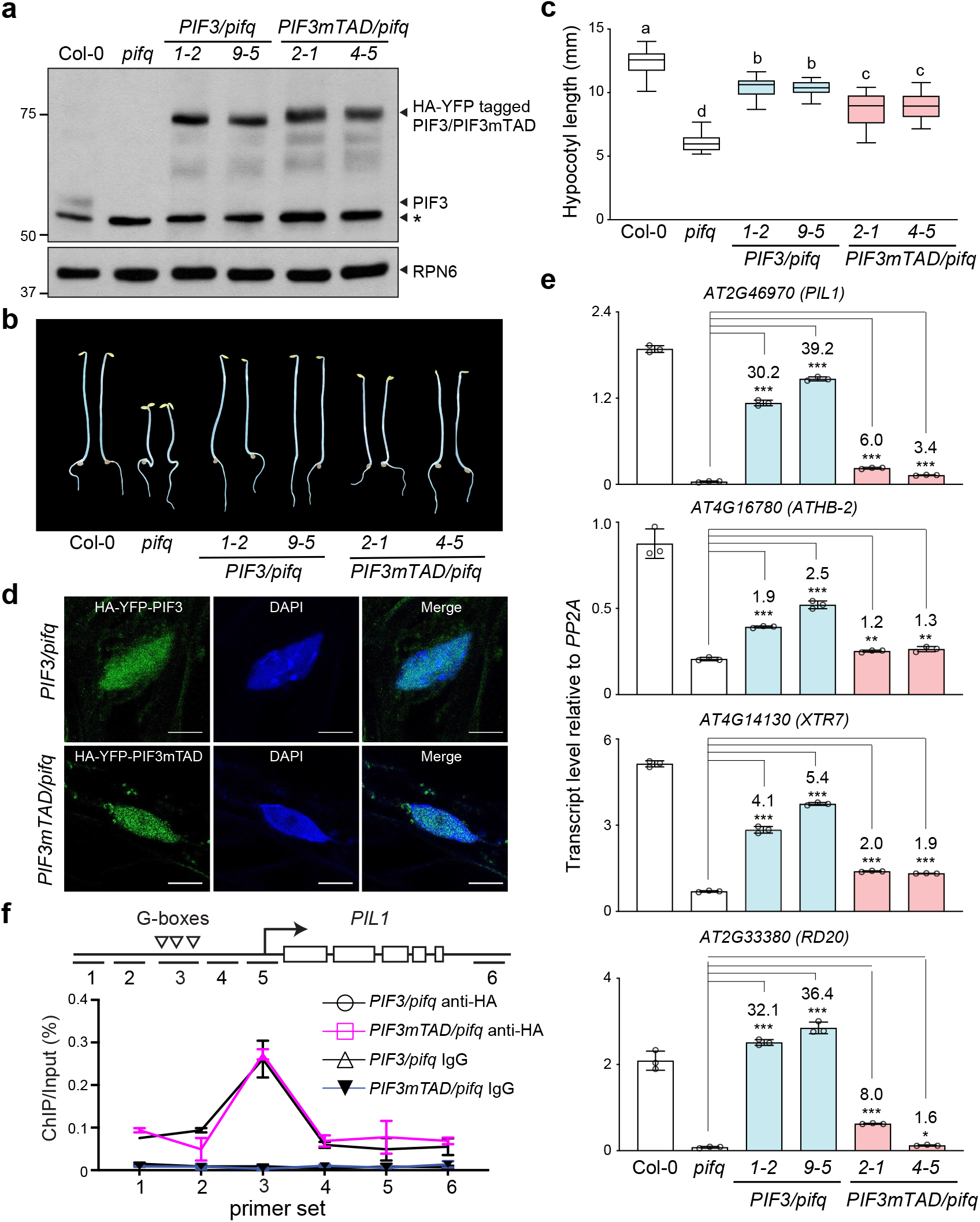
PIF3mTAD compromises PIF3’s functions in skotomorphogenesis. **a** Immunoblots showing the steady-state levels of HA-YFP-PIF3 and HA-YFP-PIF3mTAD in dark-grown seedlings of *PIF3/pifq* (*1-2* and *9-5*) and *PIF3mTAD/pifq* (*2-1* and *4-5*) transgenic lines using anti-PIF3 antibodies. Col-0 and *pifq* were used as positive and negative controls, respectively. RPN6 was used as a loading control. **b** Images of representative 4-d-old dark-grown seedlings of Col-0, *pifq*, and the transgenic *PIF3/pifq* and *PIF3mTAD/pifq* lines. **c** Hypocotyl length measurements of the lines shown in **a**. The boxes represent from the 25th to 75th percentile; the bars are equal to the median values. Samples labeled with different letters exhibited statistically significant differences in hypocotyl length (ANOVA, Tukey’s HSD, *p* < 0.05, n > 30). **d** Confocal images showing the nuclear localization of HA-YFP-PIF3 and HA-YFP-PIF3mTAD in hypocotyl epidermal cells of 4-d-old dark-grown *PIF3/pifq* (line *1-2*) and *PIF3mTAD/pifq* (line *2-1*) seedlings. Nuclei were stained with DAPI. Scale bars represent 5 μm. **e** qRT-PCR results showing the steady-state transcript levels of select PIF3 target genes in 4-d-old dark-grown seedlings of Col-0, *pifq*, and the transgenic *PIF3/pifq* and *PIF3mTAD/pifq* lines. The transcript levels were calculated relative to those of *PP2A*. Error bars represent the s.d. of three biological replicates. Numbers indicate fold changes relative to *pifq*; the statistical significance was analyzed using Student’s t-test (* *p* ≤ 0.05, ** *p* ≤ 0.01, *** *p* ≤ 0.001). **f** Chromatin immunoprecipitation (ChIP) assays showing the binding of HA-YFP-PIF3 and HA-YFP-PIF3mTAD to the G-box elements in the enhancer region of *PIL1*, as illustrated in the schematics. The ChIP experiments were performed using 4-d-old dark-grown *PIF3/pifq* (line *1-2*) and *PIF3mTAD/pifq* (line *2-1*) seedlings with anti-HA antibodies. Immunoprecipitated DNA was quantified via real-time PCR using primer sets 1 to 6 located at the *PIL1* locus. ChIP with rabbit IgG was used as a negative control. Error bars represent the s.d. of three biological replicates.

### PIF3 TAD induces *ELIP2* activation by light

PIF3 activates not only light-repressed genes but also, surprisingly, some light-inducible genes when dark-grown seedlings are exposed to light for the first time. For example, PIF3 participates in the light-dependent induction of *ELIP2* (*EARLY LIGHT-INDUCED PROTEIN 2*), which encodes a special chlorophyll binding protein that protects the plastids during initial exposure to light radiation^52^. However, it remains unclear whether *ELIP2* induction relies on PIF3’s intrinsic TAD activity or the activity of other PIF3-associated transcription factors^49^. The PIF3mTAD mutant therefore provides an opportunity to clarify the role of PIF3’s intrinsic activity in *ELIP2* induction. To that end, first, we examined the involvement of the PIF3 TAD in the early light-induced dynamics of PIF3 subnuclear distribution, phosphorylation, and abundance. During the dark-to-light transition, PHYA and PHYB recruit PIF3 to many small nuclear bodies and promote PIF3’s phosphorylation and subsequent degradation^53, 54^. These early light signaling events depend on the interactions between PIF3 and PHYs^49, 54^. Because PIF3mTAD maintains its binding to active PHYA and PHYB (Fig. 3c), as expected, HA-YFP-PIF3mTAD showed the same functionalities as HA-YFP-PIF3 in light-dependent nuclear body localization, phosphorylation, and degradation (Fig. 5a, b). However, in striking contrast, the *PIF3mTAD/pifq* lines failed to activate *ELIP2* as effectively as did the *PIF3/pifq* lines (Fig. 5c). Within one hour of R light exposure, the *PIF3/pifq* lines elevated the transcript level of *ELIP2* by more than 100 fold, whereas the transcript level of *ELIP2* increased 4-to-5-fold less in the *PIF3mTAD/pifq* lines (Fig. 5c). These results indicate that *ELIP2* activation is mainly induced by PIF3’s intrinsic TAD activity. Notably, *ELIP2* transcripts were also elevated significantly by light in *pifq,* albeit to a lesser extent (Fig. 5c), suggesting the involvement of other transcription factors besides the four PIFs in *ELIP2* induction. Therefore, PIF3 may promote the full induction of *ELIP2* through both its own TAD as well as the TADs of its associated transcription factors^49^, which provides an explanation for the remaining light response of *ELIP2* in the *PIF3mTAD/pifq* lines (Fig. 5c).

**Fig. 5.**
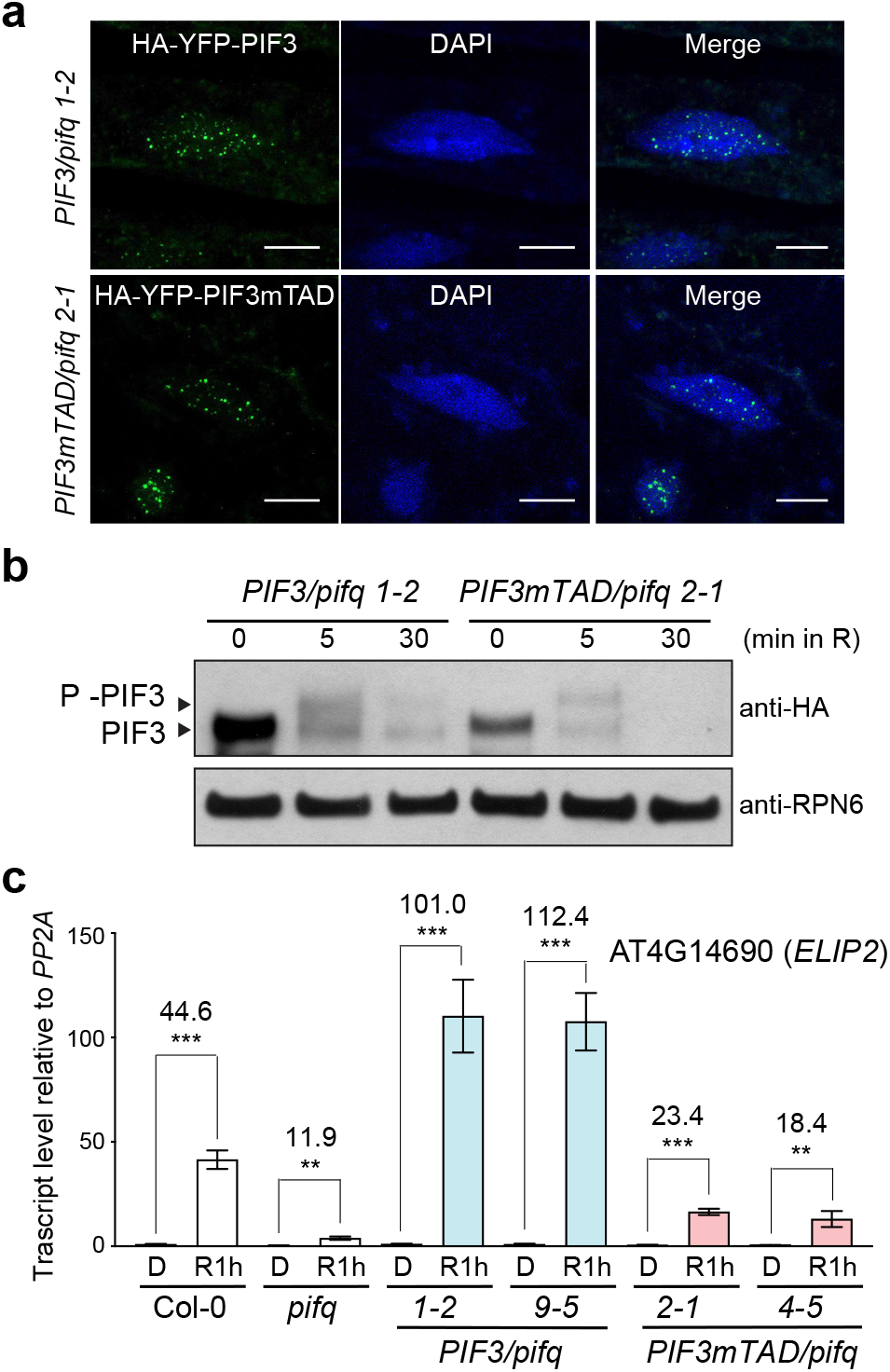
Characterization of PIF3mTAD in early light signaling. **a** Confocal images showing the nuclear body localization of HA-YFP-PIF3 and HA-YFP-PIF3mTAD in the hypocotyl epidermal cells of dark-grown *PIF3/pifq* (line *1-2*) and *PIF3mTAD/pifq* (line *2-1*) seedlings after 1 min of exposure to 10 μmol m^-2^ s^-1^ R light. Nuclei were stained with DAPI. The scale bars represent 5 μm. **b** Immunoblots showing the phosphorylation and degradation of HA-YFP-PIF3 and HA-YFP-PIF3mTAD in 4-d-old *PIF3/pifq* and *PIF3mTAD/pifq* seedlings during the dark-to-light transition at the indicated time points in 10 μ mol m^-2^ s^-1^ R light. HA-YFP-PIF3 and HA-YFP-PIF3mTAD were detected using anti-HA antibodies. RPN6 was used as a loading control. **c** qRT-PCR results showing the steady-state transcript levels of *ELIP2* in seedlings of 4-d-old dark-grown Col-0, *pifq*, and the transgenic *PIF3/pifq* and *PIF3mTAD/pifq* lines before (D) and 1 h after (R1h) exposure to 10μ mol m^-2^ s^-1^ R light. The transcript levels were calculated relative to those of *PP2A*. The numbers represent fold changes of the transcript level of *ELIP2* in R1h compared to that in D. Error bars represent the s.d. of three biological replicates.

### PIF3 TAD participates in gene activation by shade

PIF3 plays an important role in the rapid gene activation by shade or FR light, which inactivates PHYB^55^. To assess the function of the PIF3 TAD in the activation of shade-inducible genes during the light-to-shade transition, first, we characterized *PIF3/pifq* and *PIF3mTAD/pifq* lines grown in continuous R light. Interestingly, while *pifq* shows a short-hypocotyl phenotype in the light, the slow growth of *pifq* was fully rescued in *PIF3/pifq* (Fig. 6a, b). Under the R light conditions in our experiments, endogenous PIF3 did not accumulate to a detectable level. However, similar to the previously reported *PIF3* transgenic lines^27^, the two *PIF3/pifq* lines accumulated a significant amount of HA-YFP-PIF3 in the light (Fig. 6c), providing an explanation for their enhanced hypocotyl elongation in the light. On the other hand, the *PIF3mTAD/pifq* lines only partially reversed the dwarf phenotype of *pifq*, and they remained significantly shorter than the *PIF3/pifq* lines (Fig. 6a, b). HA-YFP-PIF3mTAD accumulated to similar levels as HA-YFP-PIF3 (Fig. 6c), but the transcript levels of three PIF3-target genes, *PIL1*, *ATHB-2*, and *IAA29*, in *PIF3mTAD/pifq* were similar to those in *pifq* and failed to increase to the levels in Col-0 and *PIF3/pifq* under R light conditions (Fig. 6d and Supplementary Fig. 3). Next, we treated the R-light-grown seedlings with 1 h of FR light (simulated shade) and then examined the expression of the same three PIF target genes before and after the FR treatment. The *PIF3/pifq* lines were able to dramatically enhance the expression of PIF3 target genes as Col-0 (Fig. 6d). However, similar to *pifq*, the *PIF3mTAD/pifq* lines failed to activate PIF3 target genes in FR light (Fig. 6d), confirming that the PIF3 TAD is required for gene activation by shade.

**Fig. 6.**
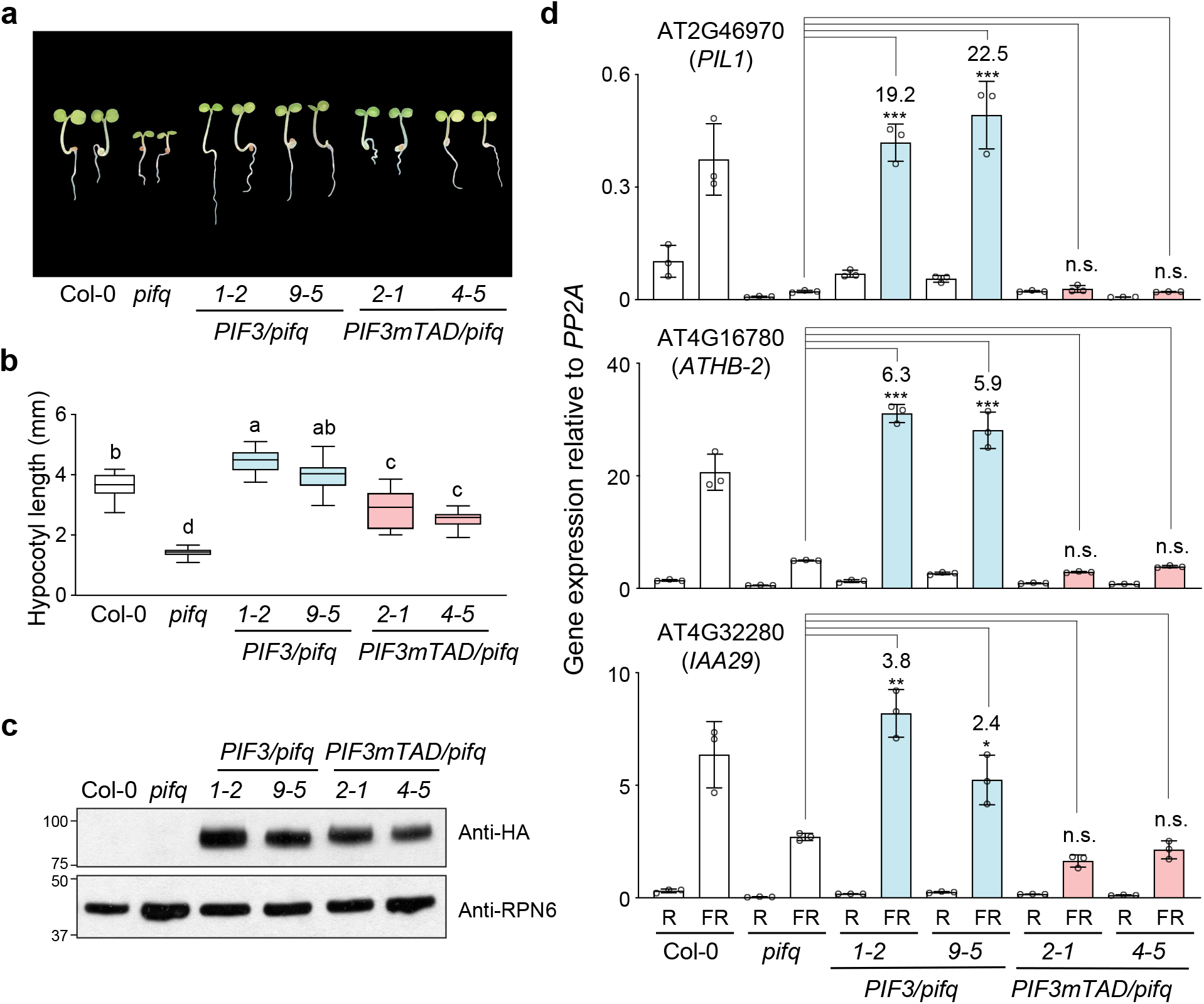
PIF3’s TAD participates in gene activation by shade. **a** Images of representative 4-d-old seedlings of Col-0, *pifq*, two *PIF3/pifq* lines, and two *PIF3mTAD/pifq* lines grown in 10μ mol m^-2^ s^-1^ R light. **b** Hypocotyl length measurements of the lines shown in **a**. The boxes represent the 25th to 75th percentiles; the bars are equal to the median values. Samples labeled with different letters exhibited statistically significant differences in hypocotyl length (ANOVA, Tukey’s HSD, *p* < 0.05, n > 30). **c** Immunoblots showing the steady-state levels of HA-YFP-PIF3 or HA-YFP-PIF3mTAD in the *PIF3/pifq* and *PIF3mTAD/pifq* seedlings shown in **a** using anti-HA antibodies. Col-0 and *pifq* were used as controls. RPN6 was used as a loading control. **d** qRT-PCR results showing the steady-state transcript levels of select PIF3 target genes in 4-d-old R-light-grown seedlings of Col-0, *pifq*, and the *PIF3/pifq* and *PIF3mTAD/pifq* lines before (R) and after a 1-h FR treatment (FR). The transcript levels were calculated relative to those of *PP2A*. Error bars represent the s.d. of three biological replicates. Numbers indicate fold changes relative to *pifq*; the the statistical significance was analyzed using Student’s t-test (* *p* ≤ 0.05, ** *p* ≤ 0.01, *** *p* ≤ 0.001); n.s. indicates the difference was either less than 2-fold or not statistically significant.

### Binding of PHYB to APB inhibits the transactivation activity of PIF3

In the shade-treatment experiments, we were intrigued by the fact that, in the *PIF3/pifq* lines, although HA-YFP-PIF3 accumulated to high levels in R light (Fig. 6c), the transcript levels of the PIF3 target genes remained low (Fig. 6d). These observations suggested that the transactivation activity of HA-YFP-PIF3 must be repressed by PHYB in the light and that this transcriptional repression is then released upon PHYB inactivation by FR light. It has been postulated that PHYB inhibits PIF3 binding to the promoters of target genes^34, 36^. However, it remains ambiguous whether PHYB influences the DNA-binding activity of PIF3 directly or indirectly through the regulation of PIF3’s TAD activity. The *PIF3mTAD/pifq* lines provide an opportunity to dissect the roles of PHYB in the regulation of PIF3 DNA-binding and transactivation activity. Because the PIF3 TAD resides in close proximity to the APB motif, we hypothesized that binding of PHYB to the APB could directly inhibit the activity of the PIF3 TAD independent of PIF3 DNA-binding activity. To discern the TAD activity of PIF3 from its DNA binding, we went back to the yeast transactivation assay and examined the activity of the Gal4 DBD-fused PIF3-N4 fragment, which includes the PIF3 TAD and the APB motif but not the bHLH DNA-binding domain (Fig. 7a). We coexpressed PIF3-N4 with PHYB conjugated with a phycocyanobilin (PCB) chromophore and performed yeast liquid β-galactosidase assays in either R or FR light to test whether the activity of PIF3-N4 could be influenced by the Pfr or the Pr form of PHYB, respectively (Fig. 7a). We found that the Pfr form of PHYB reduced the activity of PIF3-N4 by more than 70%, whereas the Pr form imposed little effect on PIF3-N4’s activity (Fig. 7b). Because PIF3 interacts with both PHYB’s N- and C-terminal modules^56^, we then tested which interaction is responsible for the transcriptional inhibition. For the N-terminal module of PHYB, we adopted a previously demonstrated biologically active form called NGB, which combines the N-terminal module of PHYB with GFP and GUS as a dimerization domain^11^. Coexpressing NGB inhibited the transactivation activity of PIF3-N4 in a light-dependent manner; in contrast, coexpressing the C-terminal module alone had little effect (Fig. 7b). These results demonstrate that the N-terminal photosensory module of PHYB can inhibit the transactivation activity of PIF3’s TAD independent of PIF3’s DNA-binding activity.

**Fig. 7.**
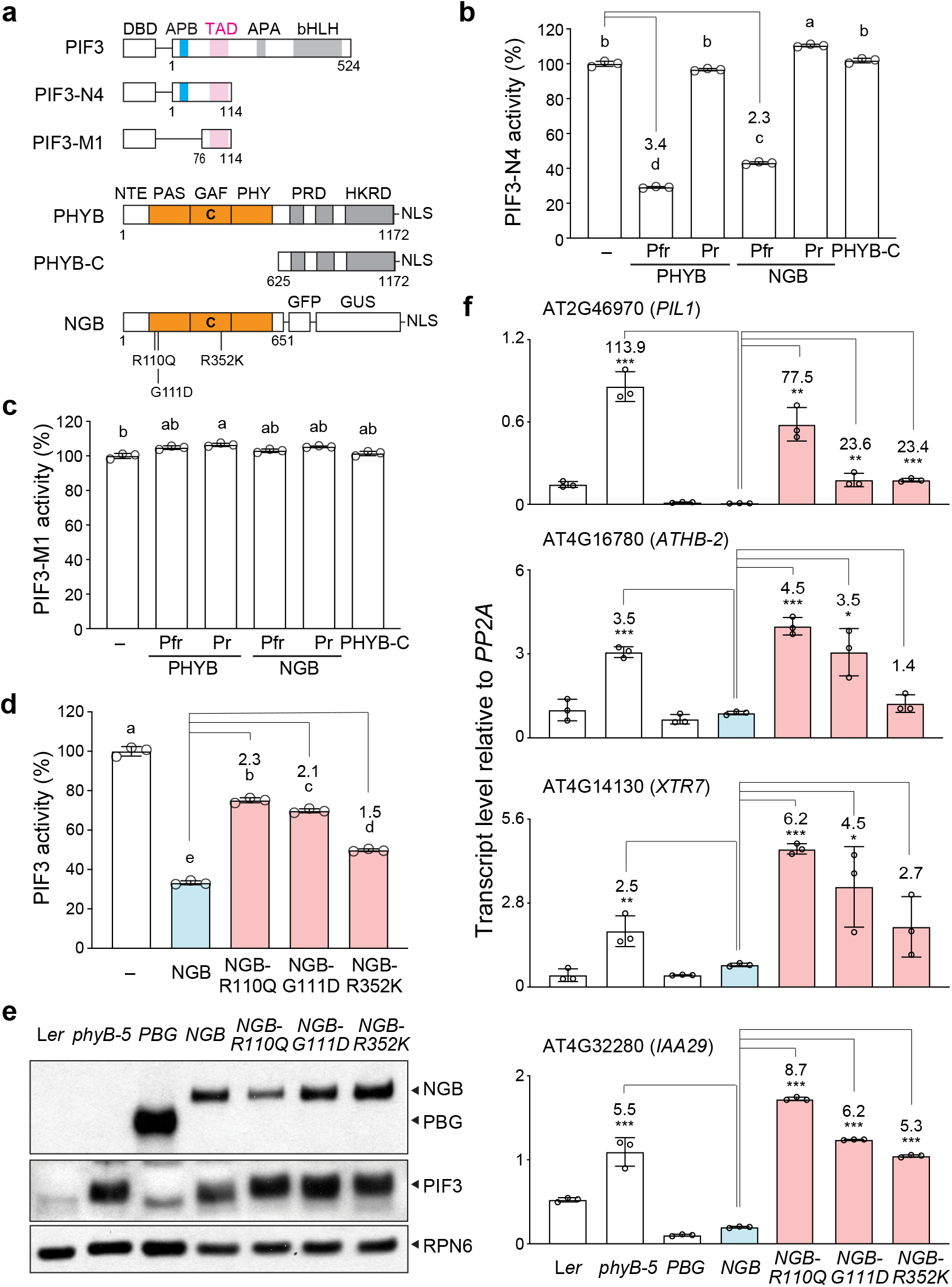
Binding of PHYB to the APB motif inhibits PIF3’s TAD activity. **a** Schematics of the PIF3 and PHYB constructs used to determine the effect of PHYB binding on PIF3’s TAD activity. NLS, nuclear localization signal; C, the cysteine residue linked to the chromophore. **b** Results of yeast liquid β-galactosidase assays showing the activity of PIF3-N4 alone (-) or coexpressed with PHYB, NGB, or PHYB-C. **c** Results of yeast liquid β-galactosidase assays showing the activity of PIF3-M1 alone (-) or coexpressed with PHYB, NGB, or PHYB-C. For **b** and **c**, the assays for PHYB and NGB were performed in either R or FR light to maintain PHYB and NGB in either the Pfr or the Pr form, respectively. The transactivation activities were calculated relative to that of the yeast strains containing the bait construct alone. Error bars represent the s.d. of three biological replicates. **d** Results of yeast liquid β-galactosidase assays showing the activity of PIF3 alone (-) or coexpressed with NGB or NGB mutants. The transactivation activities were calculated relative to that of the yeast strains expressing PIF3 alone. Samples labeled with different letters exhibited statistically significant differences in hypocotyl length (ANOVA, Tukey’s HSD, *p* < 0.05, n = 3). The numbers represent fold-changes between the NGB mutants and NGB. Error bars represent the s.d. of three biological replicates. **e** Immunoblots showing the steady-state levels of PBG, NGB, and PIF3 in 4-d-old L*er*, *phyB-5*, *PBG*, *NGB*, *NGB-R110Q*, *NGB-G111D*, and *NGB-R352K* seedlings grown in 10 μmol m^-2^s^-1^ R light. PBG and NGB were detected using anti-GFP antibodies, and PIF3 was detected using anti-PIF3 antibodies. RPN6 was used as a loading control. **f** qRT-PCR results showing the steady-state transcript levels of select PIF3 target genes in 4-d-old L*er*, *phyB-5*, *PBG*, *NGB*, *NGB-R110Q*, *NGB-G111D*, and *NGB-R352K* seedlings grown in 10 μmol m^-2^ s^-1^ R light. The transcript levels were calculated relative to those of *PP2A*. Numbers indicate fold changes relative to *NBG*. Error bars represent the s.d. of three biological replicates.

PHYB and NGB failed to repress the activity of PIF3-M1, which contains the PIF3 TAD but not the APB motif (Fig. 7c), supporting the notion that the photoinhibition of the PIF3 TAD is mediated by a direct interaction between PHYB and the APB motif. Further supporting this conclusion, three mutations in PHYB’s light-sensing knot, R110Q, G111D, and R352K, which have been previously shown to disrupt the PHYB-APB interaction^57, 58^, significantly reduced NGB’s inhibitory function on PIF3’s TAD activity (Fig. 7d). It has been previously shown that NGB can rescue the long-hypocotyl phenotype of the null *phyB-5* mutant^11^. The *in vivo* activity of NGB relies on its interaction with the APB motif in PIFs, as NGB-R110Q, NGB-G111D, and NGB-R352K failed to rescue *phyB-5*^57, 58^. Because NGB has little activity in promoting PIF3 degradation and PIF3 accumulates to high levels in light-grown *NGB* seedlings, it was proposed that NGB regulates hypocotyl growth by inhibiting the function of PIF3^10, 34–36^. Our new findings raised the possibility that NGB controls hypocotyl growth by directly repressing the TAD activity of PIF3. To further test this hypothesis, we grew *NGB, NGB-R110Q*, *NGB-G111D*, and *NGB-R352K* seedlings in continuous R light and compared the protein levels of PIF3, as well as the transcript levels of select PIF3 target genes. PIF3 accumulated to similar levels in continuous R light in *NGB*, *NGB-R110Q*, *NGB-G111D*, *NGB-R352K*, and *phyB-5*, confirming that NGB, as well as the NGB mutants, have little activity in promoting PIF3 degradation in the light (Fig. 7e)^10, 34–36^. However, despite the similar levels of PIF3, the transcript levels of PIF3 target genes in *NGB* remained low and were similar to those in *PBG*, in which PIF3 was undetectable (Fig. 7e, f), suggesting that NGB could almost fully block the activity of PIF3 *in vivo*. The transcriptional repression activity of NGB was greatly compromised by the R110Q, G111D, and R352K mutations. The transcript levels of the PIF3 target genes in *NGB-R110Q*, *NGB-G111D*, and *NGB-R352K* were elevated to levels similar to those in *phyB-5* (Fig. 6f), indicating that disrupting the NGB-APB interaction can almost completely abolish the transcriptional repression activity of NGB. These results thus provide genetic evidence supporting the model that the interaction between the light-sensing knot of PHYB’s N-terminal photosensory module to the APB motif in PIF3 directly represses the transactivation activity of the PIF3 TAD *in vivo*.

## DISCUSSION

The PHYB-PIF signaling module is at the center of plant-environment interactions that control the expression of hundreds of light-responsive genes, thereby profoundly influencing almost all aspects of plant development, growth, metabolism, and immunity^16, 20^. However, the TAD sequences of PIFs and the regulation of their intrinsic TAD activities had not been precisely determined. This study defines a 24-amino-acid TAD in PIF3 and some PIF3 paralogs, and reveals the PIF3 TAD’s conserved functional attributes of an essential ΦxxΦΦ activator motif, as well as surrounding acidic residues, which strongly resemble the TADs of the mammalian tumor suppressor p53 and the yeast activator Gcn4^40, 41, 43, 44^, indicating the conservation of sequence-specific TADs across the animal, fungal, and plant kingdoms. Moreover, we demonstrate that the light-dependent binding of the light-sensing knot of PHYB to the APB motif in PIF3 inhibits the activity of the PIF3 TAD (Fig. 8), unveiling a direct function for PHYB in the photoregulation of the transactivation activity of PIFs. These results, combined with those of previously published studies on the structure-function relationship of PHYB^10, 34–36^, indicate that the transactivation activity and stability of PIF3 are controlled by structurally separable mechanisms via PHYB’s N-terminal photosensory and C-terminal output modules, respectively (Fig. 8). While the interaction between the C-terminal module of PHYB and PIF3 promotes PIF3 degradation, the light-reversible interaction between the N-terminal photosensory module of PHYB and the APB of PIF3 directly masks PIF3’s transactivation activity (Fig. 8).

**Fig. 8.**
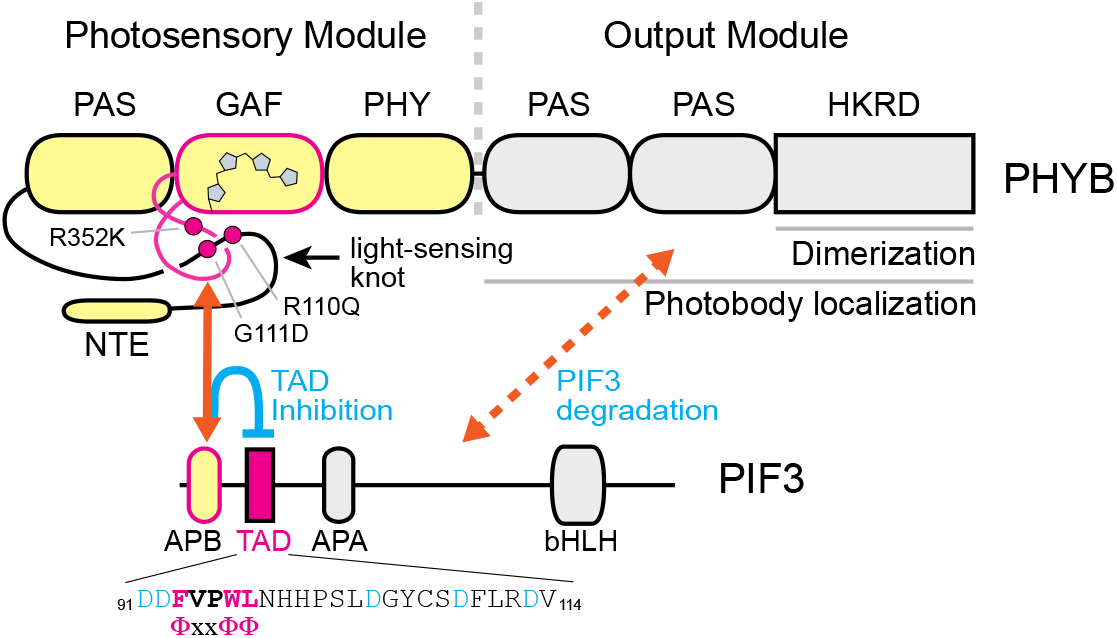
PHYB controls the stability and transactivation activity of PIF3 through structurally separable actions. The activity and stability of PIF3 are regulated separately by PHYB’s N-terminal photosensory module and C-terminal output module, respectively. Photoactivation of the N-terminal photosensory module induces the interaction between the knot lasso in PHYB and APB in PIF3, thereby masking the transactivation activity of the adjacent PIF3 TAD. In parallel, the C-terminal output module interacts weakly with PIF3 in a light-independent manner to promote PIF3 degradation. This action requires the dimerization of the C-terminal module, which is mediated by the HKRD. The orange lines represent the interactions between PHYB and PIF3. The interface involved in the interaction between PIF3 and the C-terminal module of PHYB has not been determined; therefore, a dotted line is used. The schematic of PHYB domain structure is modified from Burgie and Viestra^6^.

### PIF3 possesses a single p53-like TAD

We demonstrate that PIF3 possesses a single TAD in the aa_91-114_ region between the APB and APA motifs. The PIF3 TAD defined by this study is consistent with the previously identified activating region in yeast between amino acids 90 and 120^42^. We further narrowed down the TAD region and, more importantly, (1) showed that this is the only TAD in PIF3, (2) revealed its similarities to the TADs of p53 and Gcn4, (3) validated the TAD *in vivo*, and (4) uncovered the directly regulation of the PIF3 TAD by PHYB. The conclusion that PIF3 possesses a single p53-like TAD is solidly supported by both transactivation assays in yeast and the characterization of the PIF3mTAD mutant *in planta*. First, truncation analysis and alanine-scanning mutagenesis of PIF3 demonstrated that aa_91-114_ is the only region necessary and sufficient for PIF3’s TAD activity in yeast (Fig. 1 and Supplementary Fig. 1). Different from the prior work, which suggested a second TAD might exist between amino acids 27 and 43, our results show that aa_27-43_ is neither sufficient nor required for PIF3’s TAD activity (Fig. 1 and Supplementary Fig. 1).

Second, aa_91-114_ shares striking sequence features with the TADs of p53 and Gcn4, including an ΦxxΦΦ activator motif and flanking acidic residues. Both attributes of the PIF3 TAD are highly conserved in PIF3 orthologs in eudicots (Fig. 2a), indicating evolutionarily-conserved important functional roles. Indeed, the PIF3mTAD mutant in which the ΦxxΦΦ motif is replaced with alanines compromises PIF3’s TAD activity in yeast (Fig. 2c) as well as in the transactivation functions of PIF3 in dark, light, and shade conditions in *Arabidopsis* (Figs. 4, 5, and 6), providing strong evidence validating the function of the PIF3 TAD *in vivo*.

The TADs of p53 and Gcn4 activate transcription by binding to diverse coactivators involved in histone modification, chromatin remodeling, and the assembly of the general transcription factors and Pol II to form the so-called preinitiation complex. The p53 TADs recruit the histone acetyltransferases CBP and its paralog p300 to synergistically activate transcription^59, 60^. The Gcn4 TAD binds to the Mediator subunit Gal11/Med14^39^. Although interacting with distinct coactivators, in both cases, the intrinsically disordered TADs fold into stable amphipathic helices upon binding to their partners, and the bulky hydrophobic residues of xxΦΦ transcription activator motif interact directly with multiple residues within a shallow hydrophobic groove of the transcriptional coactivators^40, 43^. The activation mechanisms of p53 and Gcn4 inspire the hypothesis that the PIF3 TAD activates transcription via a similar mechanism. CBP/p300 and MED14 are conserved in plants. *Arabidopsis* encodes five CBP/p300-like proteins: HAC1, HAC2, HAC4, HAC5, and HAC12^61^. HACs have been shown to participate in the regulation of flowering and plant immunity^62–64^. MED14 is involved in the activation of genes in heat and cold, as well as in defense responses^65–67^. However, direct links connecting PIFs to the HACs and MED14 have yet to be revealed. It is equally possible that the p53-like TAD in PIF3 binds completely different partners in plants. A TAD does not necessarily activate transcription in different species by binding to the same coactivator. For example, VP16 exerts its functions in yeast and mammalian cells through distinct coactivators^68^. One possible PIF3-associated coactivator in plants is LEUNIG_HOMOLOG (LUH), a Groucho family protein that interacts with PIF1 and coregulates both PIF1-activated and PIF1-repressed genes^69^. Interestingly, LUH is associated with MED25 and HAC1 to facilitate gene activation by the transcription factor MYC2 in jasmonate signaling^70^, suggesting a possible link between PIF3 and both LUH and HAC1.

### Sequence-specific TADs are conserved in eukaryotes

It was demonstrated more than thirty years ago that a transcriptional activator from one species, such as the yeast activator Gal4 and the herpes simplex virus protein VP16, can act as a potent activator in animal, yeast, and plant cells, leading to the notion that the basic mechanism of transcription activation is likely conserved in eukaryotes^37, 71, 72^. However, it has been puzzling how diverse TAD sequences can carry similar functions in transcription activation across eukaryotic kingdoms. The identification of the sequence-specific ΦxxΦΦ motif in a number of animal and yeast activators implies that the mechanism of transcriptional activation could be conserved at the primary sequence level of TADs^40, 41, 43, 44^. It is important to note that the ΦxxΦΦ motif is apparently not the only activator motif as most activators in yeast lack this specific motif^40^. To our knowledge, the PIF3 TAD is the first sequence-specific TAD identified in plants. Our results therefore provide initial evidence supporting the conservation of sequence-specific TADs across the animal/fungal and plant kingdoms, suggesting that the p53/Gcn4/PIF3-type of TAD is either a result of convergent evolution or represents an ancient transactivation module that evolved before the divergence of the animal/fungal and plant lineages.

### The activity and stability of PIF3 are regulated by distinct PHYB modules

The PIF3 TAD resides between the APB and APA motifs but does not participate in PHYB or PHYA binding (Figs. 1 and 3). The APB motif mediates the light-induced interaction with the light-sensing knot of the N-terminal module of PHYB^18, 57, 58^. Our results indicate that the photoactivated N-terminal module, but not the C-terminal module, of PHYB represses the transactivation activity of PIF3 in yeast and plants (Fig. 7b). The transcriptional repression function of PHYB’s N-terminal module relies on the presence of the APB motif (Fig. 7c) and is significantly attenuated when the PHYB-APB interaction is disrupted by the R110Q, G11D, and R352K mutations in the light-sensing knot (Fig. 7d)^57, 58^. Furthermore, when PIF3 degradation is blocked in *NGB*, NGB, which contains only the N-terminal module of PHYB, is sufficient to repress the transactivation activity of PIF3 (Fig. 7e, f)^11, 34–36^. These results demonstrate that binding of the N-terminal module of PHYB to PIF3’s APB motif directly inhibits the activity of the adjacent PIF3 TAD, likely by blocking the TAD from binding to its partners. Together, the results from this study, combined with previously published data on the function of the C-terminal module in PIF3 degradation^10^, indicate that PHYB regulates the activity and stability of PIF3 through structurally separable actions of the N-terminal photosensory module and C-terminal output module, respectively (Fig. 8). Although PIF3 accumulates to very low steady-state levels in continuous light under laboratory growth conditions, it can accumulate to significant levels either temporarily in fluctuating light conditions caused by passing clouds or fluttering leaves or constantly under cold temperatures^33, 36^, we propose that the mechanism of photoinhibition of PIF3’s transactivation activity by PHYB offers an equally important mechanism, in addition to PIF3 degradation, to enable plants to rapidly finetune its responses to diverse environmental conditions.

### Relationship between PIF3 transactivation activity and DNA binding

The interaction between PHYB and PIF3 has been shown to reduce PIF3’s DNA-binding activity or enhance PIF3 sequestration away from target gene promoters independently of PHYB-mediated PIF3 degradation^34, 36^. The sequestration activity of PHYB also relies only on PHYB’s N-terminal module and is abolished by the G111D mutation in the light-sensing knot^34, 36^. Our results here thus raised the question of the relationship between PIF3’s transactivation activity and DNA binding. To discern PHYB’s function in inhibiting the PIF3 TAD from its role in regulating PIF3 DNA-binding activity, we chose to use the PIF3-N4 fragment to assess the photoinhibition of PIF3’s TAD by PHYB, because PIF3-N4 does not contain the bHLH domain, and therefore, its DNA binding relies solely on the fused Gal4-DBD in the yeast transactivation assays (Fig. 7a, b). These experiments demonstrate that the N-terminal module of PHYB can directly inhibit the transactivation activity PIF3, independently of PIF3’s bHLH activity – i.e., the photoinhibition of the activity of the PIF3 TAD by PHYB is not a consequence of a reduction in PIF3’s DNA-binding activity. On the contrary, it is conceivable that blocking the activity of PIF3’s TAD by PHYB may disrupt the interactions of PIF3 with transcriptional coactivators, thereby reducing PIF3’s association with target gene promoters. Alternatively, it is equally possible that, in addition to repressing the transactivation activity, binding of PHYB to the APB motif in PIF3 also triggers conformational changes that allosterically reduce the DNA-binding activity of the bHLH domain. The current data cannot distinguish between these two models. However, it has been shown that photoactivation of PHYB enhances PHYB’s association with DNA-bound PIF3 *in vitro*, lending further support to the model that photoactivation of PHYB can inhibit PIF3’s transactivation activity at target gene promoters^21^.

In conclusion, our results demonstrate that PIF3 possesses a p53-like TAD and unveil a novel light signaling mechanism in which light-dependent binding of the N-terminal photosensory module of PHYB to the APB motif of PIF3 directly represses the transactivation activity of the PIF3 TAD. Because the PIF3 TAD is also conserved in PIF1, PIF4, PIF5, PIF7, and PIF8, we propose that the direct transcriptional repression mechanism is a major signaling mechanism to inhibit the activity of these PIF3 paralogs, especially the ones that accumulate in the light such as PIF4, PIF5, and PIF7. It will be of interest in future work to investigate the transactivation mechanisms of individual PIFs and examine the functional significance underpinning the large variations in their transactivation activities.

## METHODS

### Plant materials, growth conditions, and hypocotyl measurement

*Arabidopsis* wild-type Columbia (Col-0) and Landsberg *erecta* (L*er*), as well as the *pifq*^23^ (Col-0) and *phyB-5*^73^(L*er*) mutants, were used as controls to characterize hypocotyl growth and gene expression under various light conditions. The *PBG*, *NGB*, *NGB-R110Q*, *NGB-G111D*, and *NGB-R352K* lines have been previously reported^11, 57^. The *PIF3/pifq* (*1-2* and *9-5*) and *PIF3mTAD/pifq* (*2-1* and *4-5*) transgenic lines were generated in this study.

*Arabidopsis* seeds were surface-sterilized and plated on half-strength Murashige and Skoog (MS) medium containing Gamborg’s vitamins (Caisson Laboratories), 0.5 mM MES pH 5.7, and 0.8 % agar (w/v) (Caisson Laboratories). Seeds were stratified in the dark at 4°C for 5 days and grown at 21°C in an LED chamber (Percival Scientific, Perry, IA) in the indicated light conditions. Fluence rates of LED light were measured with an Apogee PS200 spectroradiometer (Apogee instruments Inc., Logan, UT) and SpectraWiz spectroscopy software (StellarNet, Tampa, FL). Images of representative seedlings were captured using a Leica MZ FLIII stereo microscope (Leica Microsystems Inc., Buffalo Grove, IL) and processed using Adobe Photoshop CC (Adobe Systems, Mountain View, CA). For hypocotyl measurements, seedlings were scanned with an Epson Perfection V700 photo scanner, and hypocotyl length was qualified using the NIH ImageJ software (https://imagej.nih.gov). Box-and-whisker plots of hypocotyl measurements were generated using the Prism 8 software (GraphPad Software, San Diego, CA).

### Plasmid construction and generation of transgenic lines

The PCR primers used to generate the plasmids containing the wild-type CDS sequences of *PIF*s or *PHYB* are listed in Supplementary Table 1. All *PIF3* constructs for the yeast transactivation assays were generated by subcloning either the full-length or a fragment of *PIF3* CDS into the EcoRI/SalI sites of the pBridge vector (Clontech). The *GST-PIF3* and *GST-PIF3mTAD* constructs for GST pulldown assays were generated by subcloning *PIF3* or *PIF3mTAD* into the EcoRI/XhoI sites of the pET42b vector (Promega). The *PIF3p*::*HA-YFP-PIF3* and *PIF3p::HA-YFP-PIF3mTAD* constructs were generated by subcloning 2-kb *PIF3* promoter, *3HA-YFP*, and *PIF3* or *PIF3mTAD* into the SacI/BamHI sites of the *pJHA212G-RBCSt* vector using HiFi Assembly (New England BioLabs). The vectors for examining the transactivation activities of the PIF3 paralogs were generated by subcloning their full-length CDS sequences into the EcoRI/SalI sites of pBridge vector using HiFi Assembly. The constructs for testing the function of PHYB in inhibiting the transactivation activity of PIF3 were generated by amplifying the DNA sequences encoding full-length PHYB fused with SV40 NLS, PHYB-C (PHYB amino acids 594-1172 fused with SV40 NLS), and NGB (PHYB amino acids 1-651 fused with GFP, GUS, and SV40 NLS) and subcloning into the NotI/NdeI sites in the pBridge/PIF3-N4 or pBridge/PIF3-M1 plasmid vectors.

The constructs containing point mutants of *PIF*s or *PHYB* were generated by one of the following two methods. For PIF3 alanine scanning mutants m1 to m15, the mutations were generated in pBridge-PIF3-N1 with the Q5 Site-Directed Mutagenesis Kit (New England BioLabs). The primers used to generate the m1 to m15 mutants are listed in Supplementary Table 2. For the rest of the *PIF3* and *PHYB* point mutant constructs, two new primers from either the sense or antisense strands harboring the mutations were designed and used in combination with the primers flanking the 5’ and 3’ ends of the respective CDS sequences (Supplementary Table 1) to amplify two overlapping fragments of the respective CDS sequences. The two overlapping DNA fragments were then ligated to the indicated vectors using HiFi Assembly. The pairs of primers containing the mutated nucleotide sequences are listed in Supplementary Table 2.

### Yeast transactivation assay

Cell viability assays were performed by using either Y2HGold yeast strains (Clontech) containing a pBridge bait vector or diploid yeast strains generated by mating a Y2HGold strain containing a bait vector with a Y187 strain (Clontech) containing the pGADT7 vector. Overnight yeast cultures were diluted to an OD_600_ of 0.2 and serially diluted from 10^0^ to 10^-4^. Ten microliters of serial dilutions were spotted onto SD/-Trp media (Y2HGold strains) or SD/-Trp/-Leu (diploid strains) with or without 125 ng/ml AbA. The plates were incubated at 30°C, and pictures were taken on the third day after plating. Liquid β-galactosidase assays were performed as described in the Yeast Protocols Handbook (Clontech). A Y2HGold yeast strain containing a bait vector was mated with the Y187 strain containing the pGADT7 vector and selected on SD/-Trp/-Leu media. The diploid yeast cells were cultured in liquid SD/-Trp/-Leu media overnight, and the activity of β-galactosidase was measured by using ortho-nitrophenyl-β-galactoside (ONPG) as a substrate. To determine the effect of PHYB on PIF3’s transactivation activity, yeast strains containing indicated pBridge vector were grown in SD/-Trp/-Leu/-Met overnight and incubated with 20 µM phycocyanobilin for 4 h in the dark^74^. The yeast cells were then washed with SD-Leu/-Trp/Met media to remove unincorporated phycocyanobilin and incubated in a total of 5 mL of YPDA in 10 µmol m^-2^ s^-1^ of either R or FR light for 6 hr. The activity of β-galactosidase was measured by using ONPG as a substrate.

### GST pulldown

The GST pulldown assay was performed as described previously^75^. GST-PIF3 and GST-PIF3mTAD fusion proteins were expressed in *E. coli* strain BL21 (DE3). Cells were harvested via centrifugation and resuspended in E buffer containing 50 mM Tris-HCl pH 7.5, 100 mM NaCl, 1 mM EDTA, 1 mM EGTA, 1% DMSO, 2 mM DTT, and cOmplete^TM^ Protease Inhibitor Cocktail (Sigma-Aldrich). Cells were lysed via French press, and the lysate was centrifuged at 20,000×g for 20 min at 4 °C. The proteins were precipitated with 3.3 M ammonium sulfate via incubation for 4 h at 4 °C and centrifuged at 10,000×g for 20 min at 4 °C. The protein pellets were resuspended in E buffer. Insoluble proteins were further cleared by ultracentriguation at 50,000×g for 15 min at 4°C. The resulting lysates were dialyzed in E buffer overnight at 4°C. To immobilize GST fusion proteins, protein lysates were incubated with glutathione Sepharose beads (GE Healthcare) in E buffer for 2 h. Then, the beads were washed with E wash buffer containing 0.1% IGEPAL CA-630. Apoproteins of PHYA-HA and PHYB-HA were prepared using a TNT T7 Coupled Reticulocyte Lysate system (Promega) as described previously^75^. Holoproteins of PHYA-HA and PHYB-HA were generated by incubating with 20 µM PCB for 1 h in the dark. The *in vitro*-translated proteins were incubated with the immobilized GST fusion proteins in E wash buffer for 2 h at 4°C. The beads were washed with E wash buffer, the bound proteins were eluted by boiling in Laemmli sample buffer, and the samples were subjected to SDS-PAGE.

### Protein extraction and immunoblots

Total proteins from 100 mg of 4-day-old seedlings were extracted in 300 µl of extraction buffer (100 mM Tris-HCl pH 7.5, 100 mM NaCl, 5 mM EDTA pH 8.0, 5% SDS, 20% glycerol, 20 mM DTT, 40 mM β-mercaptoethanol, 2 mM PMSF, cOmplete^TM^ Protease Inhibitor Cocktail [Sigma-Aldrich], 80 µM MG132 [Sigma-Aldrich] and 80 µM MG115 [Sigma-Aldrich], 1% phosphatase inhibitor cocktail 3 [Sigma-Aldrich], 10 mM N-ethylmaleimide, and 0.01% bromophenol blue). Samples were immediately boiled for 10 min and centrifuged at 16,000×g for 10 min. Proteins were separated via SDS-PAGE and blotted onto a nitrocellulose membrane. The membrane was first probed with the indicated primary antibodies and then incubated with goat anti-rabbit (Bio-Rad, cat. no. 1706515) or goat anti-mouse (Bio-Rad, cat. no. 1706516) secondary antibodies conjugated with horseradish peroxidase at a 1:5000 dilution. The signals were detected via a chemiluminescence reaction using the SuperSignal West Dura Extended Duration Substrate (ThermoFisher Scientific). Rabbit polyclonal anti-PIF3^76^, goat polyclonal anti-HA (GenScript, cat. no. A00168), rabbit polyclonal anti-GFP (Abcam, cat. no. ab290), and rabbit polyclonal anti-RPN6 (Enzo Life Sciences, cat. no. BML-PW8370-0100) antibodies were used at a 1:1000 dilution.

### Confocal imaging

Confocal analysis of HA-YFP-PIF3 and HA-YFP-PIF3mTAD was performed as previously described with minor modifications^77^. Seedlings were fixed in 2% paraformaldehyde in PBS under vacuum for 15☐min and then washed three times with 50 mM NH_4_Cl in PBS for 5☐min three times. The seedlings were permeabilized with 0.2% Triton X-100 in PBS for 5☐min, incubated with 300 nM DAPI in PBS for 10 min. The seedlings were washed three times with PBS for 5 min three times, and mounted with ProLong^TM^ Diamond Antifade (ThermoFisher Scientific) on a slide. The slides were left to cure overnight in the dark, sealed with nail polish, and stored at 4°C until imaging. Nuclei of hypocotyl epidermal cells were imaged using a Zeiss Axio Observer Z1 inverted microscope equipped with a Plan-Apochromat 100×/1.4 oil-immersion objective and an Axiocam 506 mono camera (Carl Zeiss, Jena, Germany). Fluorescence was detected with the following Zeiss filter sets: YFP, exciter 500/25 nm/nm, emitter 535/40 nm/nm (Zeiss Filter Set 46); DAPI, exciter 365 nm, emitter 445/50nm/nm (Zeiss Filter Set 49). Images were collected using Zeiss ZEN software and processed using Adobe Creative Cloud (Adobe, San Jose, CA).

### RNA extraction and qRT-PCR

Total RNA was extracted from seedlings using a Quick-RNA MiniPrep kit with on-column DNase I treatment (Zymo Research). cDNA synthesis was performed with 2 µg of total RNA using oligo(dT) primers and Superscript II First-Strand cDNA Synthesis Kit (Thermo Fisher Scientific). Quantitative RT-PCR was performed with FastStart Universal SYBR Green Master Mix and a LightCycler 96 Real-Time PCR System (Roche). Transcript levels of genes were calculated relative to the level of *PP2A*. Genes and primer sets used for qRT-PCR are listed in Supplementary Table 3.

### Chromatin immunoprecipitation

ChIP assays were performed using chromatin isolated from 4-d-old dark-grown *PIF3/pifq* and *PIF3mTAD/pifq* transgenic lines. Seedlings were ground in liquid nitrogen and resuspended in nuclear isolation buffer (10 mM HEPES pH 7.6, 1 M sucrose, 5 mM KCl, 5 mM MgCl_2_, 5 mM EDTA, 14 mM β-mercaptoethanol, protease inhibitor cocktail, 40 µM MG132, and 40 µM MG115) containing 1% formaldehyde, 0.6% Triton X-100, and 0.4 mM PMSF. Samples were incubated at room temperature for 10 min for crosslinking, and then 125 mM glycine was added to terminate the crosslinking. The lysate was filtered through two-layer Miracloth, and the cleared lysate was centrifuged at 3,000×g for 10 min. The nuclei pellet was resuspended in nuclear isolation buffer, loaded on top of a 15% Percoll solution (15% Percoll, 10 mM HEPES pH 8.0, 1 M sucrose, 5 mM KCl, 5 mM MgCl_2_, 5 mM EDTA), and centrifuged at 3,000×g for 5 min. The enriched nuclear pellet was lysed with nuclear lysis buffer (50 mM Tris-HCl pH 7.5, 1% SDS, 10 mM EDTA). Then, sonication was performed using a Covaris S2 ultrasonicator (Covaris, Inc., Woburn, MA). The lysate was centrifuged at 13,000×g for 3 min at 4°C to remove any debris. The nuclear lysate was diluted with ChIP dilution buffer (15 mM Tris-HCl pH 7.5, 150 m NaCl, 1 mM EDTA, 1% Triton X-100). The lysate was incubated with 1 µg of rabbit polyclonal anti-HA (Abcam, cat. no. ab9110) or rabbit IgG (Cell Signaling Technology, cat. no. 2729S) for 3 h at 4 °C. Immunoprecipitated chromatin with anti-HA antibody or IgG was then incubated with Dynabeads Protein G (Thermo Fisher Scientific, cat. no. 10003D) for 2 h. The beads were washed with ChIP dilution buffer, low-salt wash buffer (20 mM Tris-HCl pH 8.0, 150 mM NaCl, 0.1% SDS, 1% Triton X-100, 2 mM EDTA), high-salt wash buffer (20 mM Tris-HCl pH 8.0, 500 mM NaCl, 0.1% SDS, 1% Triton X-100, 2 mM EDTA), LiCl wash buffer (10 mM Tris-HCl pH 8.0, 0.25 M LiCl, 1% IGEPAL CA-630, 1% sodium deoxycholate, 1 mM EDTA), and TE buffer (10 mM Tris-HCl pH 8.0, 1 mM EDTA). The chromatin was eluted from the beads with elution buffer (1% SDS, 0.1 M NaHCO_3_). Reverse cross-linking was performed by adding 20 µl of 5 M NaCl to the eluates and incubating at 65 °C overnight. Then, the immunoprecipitated proteins were digested by adding 10 µl of 0.5 M EDTA pH 8.0, 20 µl of 1M Tris-HCl pH 6.5 and 20 µg proteinase K and incubating at 50°C for 2 h. The final chromatin was purified with ChIP DNA Clean & Concentrator (Zymo Research). Primer sets used for ChIP-qPCR analysis are listed in Supplementary Table 4.

## DATA AVAILABILITY

*Arabidopsis* mutants and transgenic lines, as well as plasmids generated during the current study, are available from the corresponding author upon reasonable request.

## ACKNOWLEDGEMENTS

We thank Dr. Elise Pasoreck for providing valuable comments and suggestions on the manuscript. This work was supported by National Institute of General Medical Sciences grants R01GM087388 and R01GM132765 to M.C. and National Institute of Food and Agriculture hatch projects CA-R-BPS-5186-H and CA-R-BPS-5084-H to M.C. and X.C., respectively. Q.S. was supported by Guangdong Innovation Team Project 2014ZT05S078.

## AUTHOR CONTRIBUTIONS

C.Y.Y., Q.S., J.H., Y.Q., L.L., B.M., X.C., and M.C. conceived the original research plan; M.C. and X.C. supervised the experiments; C.Y.Y., Q.S., J.H., Y.Q., L.L., R.J.K., E.C., and J.H. performed the experiments; C.Y.Y., Q.S., J.H. Y.Q. L.L., R.J.K., E.C., J.H., N.M., P.Z., L.S., A.N., B.M., X.C., M.C. analyzed the data; A.N. provided the transgenic lines expressing NGB and NGB mutants; M.C., C.Y.Y., Q.S., and J.H. wrote the article with contributions from all authors.

## SUPPLEMENTARY INFORMATION

**Supplementary Fig. 1. The aa_1-90_ region does not exhibit transactivation activity in yeast.** A series of N-terminal fragments of PIF3 between amino acids 1 and 101 were fused with Gal4-DBD as shown in the schematics and examined for their self-activation activity in yeast. Serial dilutions of the yeast strains containing the respective constructs were grown on either SD/-Trp/+AbA or SD/-Trp (control) media.

**Supplementary Fig. 2. Abolishing the transactivation activity of PIF3 requires multiple mutations in the TAD. a** Sequence alignment of the PIF3 TAD and the mTAD1 mutant, in which the three hydrophobic residues in the ΦxxΦΦ activator motif were replaced with either an arginine or a serine (labeled in orange). The black bar indicates the region of amino acids 91 to 100, containing the conserved activator motif. **b** Yeast transactivation assays of DBD-PIF3-N1 (PIF3-N1) and DBD-PIF3-N1 mutants with individual amino acids between 91 and 100 replaced with an alanine. The yeast strains containing the respective constructs were grown on either SD/-Trp/+AbA or SD/-Trp (control) media. **c** Yeast transactivation assays of DBD-PIF3 (PIF3) and DBD-PIF3mTAD1 (PIF3mTAD1). Serial dilutions of the yeast strains containing the respective constructs were grown on either SD/-Trp/-Leu/+AbA or SD/-Trp/-Leu (control) media. Yeast strains containing p53 and either the SV40 large T-antigen (p53+T) or lamin (p53+Lam) were used as positive and negative controls, respectively.

**Supplementary Fig. 3. *PIF3mTAD/pifq* lines exhibit reduced activity in target gene expression in R light.** qRT-PCR results showing the steady-state transcript levels of select PIF3 target genes in 4-d-old R-light-grown seedlings of Col-0, *pifq*, the *PIF3/pifq* and *PIF3mTAD/pifq* lines. The transcript levels were calculated relative to those of *PP2A*. Error bars represent the s.d. of three biological replicates. Numbers indicate fold changes relative to *pifq*; the statistical significance was analyzed using Student’s t-test (** *p* ≤ 0.01, *** *p* ≤ 0.001); n.s. indicates the difference is either less than 2-fold or not statistically significant.

**Supplementary Table 1.** Primers used for making the constructs of PIF3 and PIF3 paralogs.

**Supplementary Table 2.** Primers used for generating the mutant constructs of PIFs and PHYB.

**Supplementary Table 3.** Primers for qRT-PCR analyses.

**Supplementary Table 4.** Primer sets for ChIP-qPCR analysis of the *PIL1* locus.

