## Supplementary Information for "Direct photoresponsive inhibition of a p53-like transcription activation domain in PIF3 by *Arabidopsis* phytochrome B"

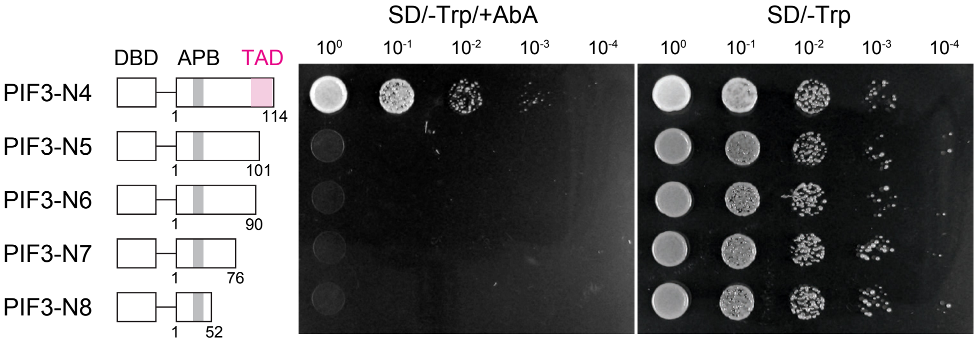


**Supplementary Fig. 1. The aa$_{1\text{-}90}$ region does not exhibit transactivation activity in yeast.** A series of N-terminal fragments of PIF3 between amino acids 1 and 101 were fused with Gal4-DBD as shown in the schematics and examined for their self-activation activity in yeast. Serial dilutions of the yeast strains containing the respective constructs were grown on either SD/-Trp/+AbA or SD/-Trp (control) media.

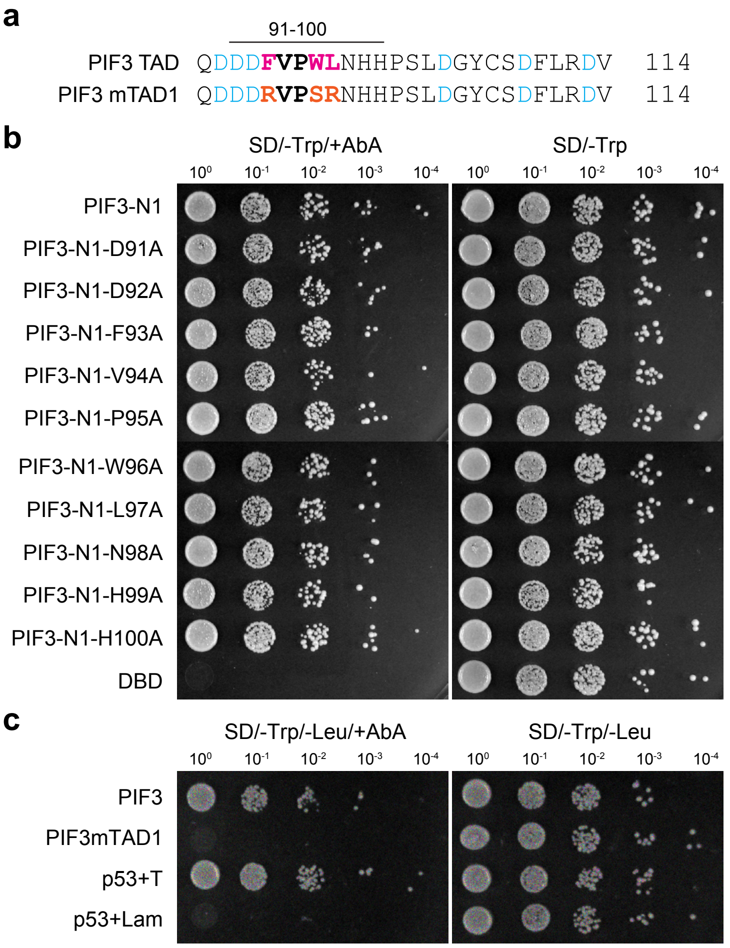


**Suplementary Fig. 2. Abolishing the transactivation activity of PIF3 requires multiple mutations in the TAD. a** Sequence alignment of the PIF3 TAD and the mTAD1 mutant, in which the three hydrophobic residues in the ΦxxΦΦ activator motif were replaced with either an arginine or a serine (labeled in orange). The black bar indicates the region of amino acids 91 to 100, containing the conserved activator motif. **b** Yeast transactivation assays of DBD-PIF3-N1 (PIF3-N1) and DBD-PIF3-N1 mutants with individual amino acids between 91 and 100 replaced with an alanine. The yeast strains containing the respective constructs were grown on either SD/-Trp/+AbA or SD/-Trp (control) media. **c** Yeast transactivation assays of DBD-PIF3 (PIF3) and DBD-PIF3mTAD1 (PIF3mTAD1). Serial dilutions of the yeast strains containing the respective constructs were grown on either SD/-Trp/-Leu/+AbA or SD/-Trp/-Leu (control) media. Yeast strains containing p53 and either the SV40 large T-antigen (p53+T) or lamin (p53+Lam) were used as positive and negative controls, respectively.

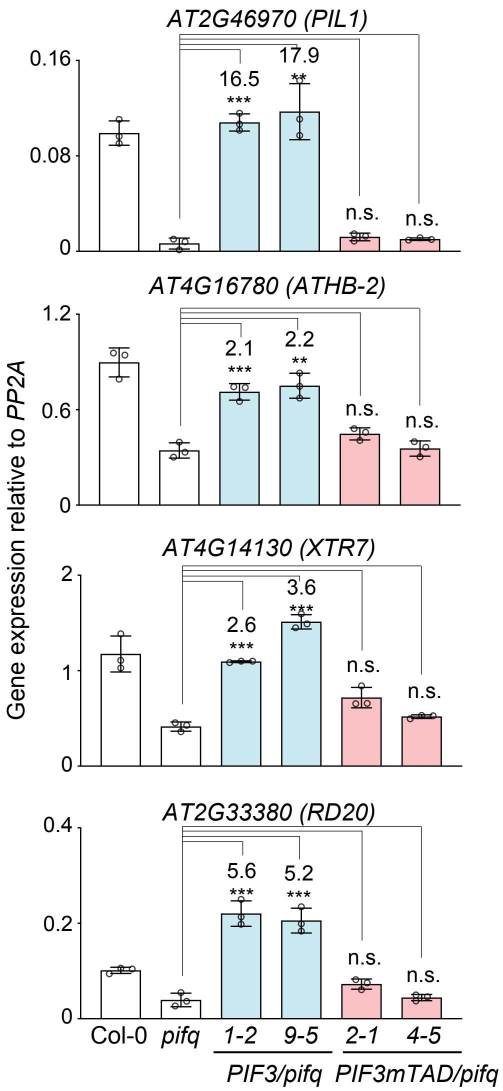


**Supplementary Fig. 3. *PIF3mTAD/pifq* lines exhibit reduced activity in target gene expression in R light.** qRT-PCR results showing the steady-state transcript levels of select PIF3 target genes in 4-d-old R-light-grown seedlings of Col-0, *pifq*, and the *PIF3/pifq* and *PIF3mTAD/pifq* lines. The transcript levels were calculated relative to those of *PP2A*. Error bars represent the s.d. of three biological replicates. Numbers indicate fold changes relative to *pifq*; the statistical significance was analyzed using Student's t-test (** $p \leq 0.01$, *** $p \leq 0.001$); n.s. indicates the difference is either less than 2-fold or not statistically significant.

**Supplementary Table 1. PCR primers used for making the constructs of PIF3 and PIF3 paralogs.**

| Plasmid name | Insert | Vector | Forward primer | Reverse primer |
|---|---|---|---|---|
| pBridge-PIF3 | *PIF3* | pBridge | acagttgactgtatcgccggctatgcctctgtttgagcttttcagg | ggaattagcttggctgcaggtcacgacgatccacaaaactg |
| pBridge-PIF3-N1 | *PIF3-N1* | pBridge | acagttgactgtatcgccggctatgcctctgtttgagcttttcagg | ggaattagcttggctgcaggtcaacctgcttcctttcttccatc |
| pBridge-PIF3-N2 | *PIF3-N2* | pBridge | acagttgactgtatcgccggctatgcctctgtttgagcttttcagg | ggaattagcttggctgcaggttacgggctttgaggactcttttc |
| pBridge-PIF3-N3 | *PIF3-N3* | pBridge | acagttgactgtatcgccggctatgcctctgtttgagcttttcagg | ggaattagcttggctgcaggttacggattggtttgttggctagg |
| pBridge-PIF3-N4 | *PIF3-N4* | pBridge | acagttgactgtatcgccggctatgcctctgtttgagcttttcagg | ggaattagcttggctgcaggttagggatgatgattcaaccatgg |
| pBridge-PIF3-N5 | *PIF3-N5* | pBridge | acagttgactgtatcgccggctatgcctctgtttgagcttttcagg | ggaattagcttggctgcaggtcagggatgatgattcaaccat |
| pBridge-PIF3-N6 | *PIF3-N6* | pBridge | acagttgactgtatcgccggctatgcctctgtttgagcttttcagg | ggaattagcttggctgcaggtcagtcttgactcaaacccgtc |
| pBridge-PIF3-N7 | *PIF3-N7* | pBridge | acagttgactgtatcgccggctatgcctctgtttgagcttttcagg | ggaattagcttggctgcaggtcagatctcgtccaccatagtc |
| pBridge-PIF3-N8 | *PIF3-N8* | pBridge | acagttgactgtatcgccggctatgcctctgtttgagcttttcagg | ggaattagcttggctgcaggtcaaggaatgttcctcgatcta |
| pBridge-PIF3-C1 | *PIF3-C1* | pBridge | acagttgactgtatcgccggctatccctatgtcagtgccatc | ggaattagcttggctgcaggtcacgacgatccacaaaactg |
| pBridge-PIF3-C2 | *PIF3-C2* | pBridge | acagttgactgtatcgccggctgatgactttgttccatggt | ggaattagcttggctgcaggtcacgacgatccacaaaactg |
| pBridge-PIF3-C3 | *PIF3-C3* | pBridge | acagttgactgtatcgccggctccctcccttgatggatatt | ggaattagcttggctgcaggtcacgacgatccacaaaactg |
| pBridge-PIF3-M1 | *PIF3-M1* | pBridge | acagttgactgtatcgccggctatccctatgtcagtgccatc | ggaattagcttggctgcaggttagggatgatgattcaaccatgg |
| pBridge-PIF3-M2 | *PIF3-M2* | pBridge | acagttgactgtatcgccggctgatgactttgttccatggt | ggaattagcttggctgcaggttagggatgatgattcaaccatgg |
| pET42b-GST-PIF3 | *PIF3* | pET42b | ggcgaattctatgcctctgtttgagcttttcag | ccgctcgagcgacgatccacaaaactgatc |
| pJHA212G-3HA-YFP-PIF3 | *PIF3p* | pJHA212 G-RBCSt | catgattacgaattcgagctataaaccagaagatgcaac | ggtgttgcgttttacagaa |
| | *3HA-YFP* | | gtaaaacgcaacaccatgtacccatacgatgttc | gaaaagctcaaacagaggcataggggtgggagttggtgt |
| | *PIF3* | | atgcctctgtttgagcttt | gcaggtcgactctagagtcacgacgatccacaaaac |
| pJHA212G-3HA-YFP-PIF3mTAD | *PIF3p* | pJHA212 G-RBCSt | catgattacgaattcgagctataaaccagaagatgcaac | ggtgttgcgttttacagaa |
| | *3HA-YFP* | | gtaaaacgcaacaccatgtacccatacgatgttc | gaaaagctcaaacagaggcataggggtgggagttggtgt |
| | *PIF3* | | atgcctctgtttgagcttt | gcaggtcgactctagagtcacgacgatccacaaaac |
| pBridge-PIF3/PHYB | *PIF3/PHYB-NLS* | pBridge | catccatacaatgggccaatggtttccggagtcggggg | agatcttcgggctaatgcttacacctttctcttcttcttaggatatggcatcatcagcat |

| | | | | |
|---|---|---|---|---|
| pBridge-PIF3/PHYB-C | *PIF3/PHYB-C-NLS* | pBridge | catccatacaatgggccaatgaactctaaagttgtcgat | agatcttcgggctaatgcttacacctttctcttcttcttaggatatggcatcatcagcat |
| pBridge-PIF3/NGB | *PIF3/NGB* | pBridge | catccatacaatgggccaatggtttccggagtcggggg | aagatcttcgggctaatgcctatcctccaacctttctc |
| pBridge-PIF1 | *PIF1* | pBridge | acagttgactgtatcgccggctatgcatcattttgtccctg | ggaattagcttggctgcaggttaacctgttgtgtggttt |
| pBridge-PIF4 | *PIF4* | pBridge | acagttgactgtatcgccggctatggaacaccaaggttgg | ggaattagcttggctgcaggctagtggtccaaacgaga |
| pBridge-PIF5 | *PIF5* | pBridge | gactgtatcgccggctatggaacaagtgtttgctga | ttagcttggctgcaggtcagcctattttacccatat |
| pBridge-PIF7 | *PIF7* | pBridge | gactgtatcgccggctatgtcgaattatggagttaa | ttagcttggctgcaggctaatctcttttctcatgat |
| pBridge-PIF8 | *PIF8* | pBridge | gactgtatcgccggctatgtcccaatgtgttccaaa | ttagcttggctgcaggtcattttggattcgaaggaggag |
| pBridge-PIL1 | *PIL1* | pBridge | ttgactgtatcgccggctatggaagcaaaacccttagc | ttagcttggctgcaggttagtttggcgagcgataat |
| pBridge-PIL2 | *PIL2* | pBridge | gactgtatcgccggctatgatgttcttaccaaccga | ttagcttggctgcaggtcatctgttagttttccttg |

**Supplementary Table 2. Primers used for generating the mutant constructs of PIFs and PHYB.**

| Plasmid name | Gene name | Vector | Forward primer | Reverse primer |
|---|---|---|---|---|
| pBridge-PIF3-N1m1 | *PIF3-N1m1* | pBridge | gagctggtgtgggaaaatggtc | ccattttcccacaccagctcagcggctgcggcagcaggtggagaagggttcctgtc |
| pBridge-PIF3-N1m2 | *PIF3-N1m2* | pBridge | aatggtcagatatcaactcaaag | tgagttgatatctgaccattagcggctgcggcagccacaacttcatctacaggtgg |
| pBridge-PIF3-N1m3 | *PIF3-N1m3* | pBridge | actcaaagtcagtcaagtagatc | ctacttgactgactttgagtagcggctgcggcagcttcccacaccagctccacaac |
| pBridge-PIF3-N1m4 | *PIF3-N1m4* | pBridge | agtagatcgaggaacattcctc | ggaatgttcctcgatctactagcggctgcggcagctgatatctgaccattttcccac |
| pBridge-PIF3-N1m5 | *PIF3-N1m5* | pBridge | attcctccaccacaagcaaac | tttgcttgtggtggaggaatagcggctgcggcagctgactgactttgagttgatatc |
| pBridge-PIF3-N1m6 | *PIF3-N1m6* | pBridge | gcaaactcttctagagctagag | ctagctctagaagagtttgcagcggctgcggcagcgttcctcgatctacttgactg |
| pBridge-PIF3-N1m7 | *PIF3-N1m7* | pBridge | gctagagagattggaaatggc | ccatttccaatctctctagcagcggctgcggcagcttgtggtggaggaatgttcctc |
| pBridge-PIF3-N1m8 | *PIF3-N1m8* | pBridge | aatggctcaaagacgactatg | atagtcgtctttgagccattagcggctgcggcagctctagaagagtttgcttgtgg |
| pBridge-PIF3-N1m9 | *PIF3-N1m9* | pBridge | actatggtggacgagatccc | gggatctcgtccaccatagtagcggctgcggcagctccaatctctctagctctagaag |
| pBridge-PIF3-N1m10 | *PIF3-N1m10* | pBridge | cttcagtggagactgatatgcc | gatggcactgacatagggatagcggctgcggcagccgtctttgagccatttccaatc |
| pBridge-PIF3-N1m11 | *PIF3-N1m11* | pBridge | ccatcactaatgacgggtttg | caaacccgtcattagtgatggagcggctgcggcagcctcgtccaccatagtcgtctttg |
| pBridge-PIF3-N1m12 | *PIF3-N1m12* | pBridge | ggtttgagtcaagacgatgac | tcatcgtcttgactcaaaccagcggctgcggcagccactgacatagggatctcgtc |
| pBridge-PIF3-N1m13 | *PIF3-N1m13* | pBridge | gatgactttgttccatggttg | aaccatggaacaaagtcatcagcggctgcggcagccgtcattagtgatggcactgac |
| pBridge-PIF3-N1m14 | *PIF3-N1m14* | pBridge | tggttgaatcatcatccctcc | gagggatgatgattcaaccaagcggctgcggcagcgtcttgactcaaacccgtcattag |
| pBridge-PIF3-N1m15 | *PIF3-N1m15* | pBridge | ccctcccttgatggatattgc | caatatccatcaagggagggagcggctgcggcagctggaacaaagtcatcgtcttg |
| pBridge-PIF3-N1m16 | *PIF3-N1m16* | pBridge | tattgctctgatttcttgcg | gcaagaaatcagagcaatatgcagcagcggcggcatgatgattcaaccatggaa |
| pBridge-PIF3-N1m17 | *PIF3-N1m17* | pBridge | ttgcgtgatgtgtcgtctc | gacgacacatcacgcaaggcagcagcggcagctccatcaagggagggatg |
| pBridge-PIF3-N1m18 | *PIF3-N1m18* | pBridge | tcgtctcctgttactgtca | gacagtaacaggagacgacgcagcagccgcgaaatcagagcaatatccatc |
| pBridge-PIF3-N1-D91A | *PIF3-N1-D91A* | pBridge | agtcaagacgctgactttgttccatggttga | aacaaagtcagcgtcttgactcaaacccg |

| | | | | |
|---|---|---|---|---|
| pBridge-PIF3-N1-D92A | *PIF3-N1-D92A* | pBridge | caagacgatgcctttgttccatggttgaatc | tggaacaaaggcatcgtcttgactcaaacc |
| pBridge-PIF3-N1-F93A | *PIF3-N1-F93A* | pBridge | agacgatgacgctgttccatggttgaatcat | ccatggaacagcgtcatcgtcttgactcaa |
| pBridge-PIF3-N1-V94A | *PIF3-N1-V94A* | pBridge | gatgactttgctccatggttgaatcatcat | caaccatggagcaaagtcatcgtcttgactc |
| pBridge-PIF3-N1-P95A | *PIF3-N1-P95A* | pBridge | tgactttgttgcatggttgaatcatcatc | attcaaccatgcaacaaagtcatcgtcttg |
| pBridge-PIF3-N1-W96A | *PIF3-N1-W96A* | pBridge | caagacgatgactttgttccagcattgaatcatcatccctcccttg | caagggagggatgatgattcaatgctggaacaaagtcatcgtcttg |
| pBridge-PIF3-N1-L97A | *PIF3-N1-L97A* | pBridge | gacgatgactttgttccatgggcaaatcatcatccctcccttgatg | catcaagggagggatgatgatttgcccatggaacaaagtcatcgtc |
| pBridge-PIF3-N1-N98A | *PIF3-N1-N98A* | pBridge | gatgactttgttccatggttggcacatcatccctcccttgatgg | ccatcaagggagggatgatgtgccaaccatggaacaaagtcatc |
| pBridge-PIF3-N1-H99A | *PIF3-N1-H99A* | pBridge | gactttgttccatggttgaatgcacatccctcccttgatggatattg | caatatccatcaagggagggatgtgcattcaaccatggaacaaagtc |
| pBridge-PIF3-N1-H100A | *PIF3-N1-H100A* | pBridge | ctttgttccatggttgaatcatgcaccctcccttgatggatattg | caatatccatcaagggagggtgcatgattcaaccatggaacaaag |
| pBridge-PIF3mTAD | *PIF3 mTAD* | pBridge | gctgctgcagcggcgaatcatcatccctccc | cgccgctgcagcagcgtcatcgtcttgactcaa |
| pBridge-PIF3/NGB-R110Q | *NGB-R110Q* | pBridge | aatccagcaaggtggttacattcagcc | tgtaaccaccttgctggattcgagagagat |
| pBridge-PIF3/NGB-G111D | *NGB-G111D* | pBridge | tccagcgagatggttacattcagcctttc | gaggagccttaagagtagaaccaaccaag |
| pBridge-PIF3/NGB-R352K | *NGB-R352K* | pBridge | tctactcttaaggctcctcatggttgtca | gaggagccttaagagtagaaccaaccaag |
| pBridge-PIF3mTAD1 | *PIF3mTAD1* | pBridge | cgtgttccatctcgtaatcatcatccctcccttgatgga | atgattacgagatggaacacggtcatcgtcttgaca |
| pBridge-PIF1mTAD | *PIF1mTAD* | pBridge | gatgaagctgcggcggctgcgcattatcctctccgt | gataatgcgcagccgccgcagcttcatcttcttgaata |
| pBridge-PIF4mTAD | *PIF4mTAD* | pBridge | atcaagaagctgcggcggctgcgcaataccctccagatgaag | gtattgcgcagccgccgcagcttcttgatcttcaagaaag |
| pBridge-PIF5mTAD | *PIF5mTAD* | pBridge | caagaagctgcggcggctgcgcaataccctccggatgacgtc | agggtattgcgcagccgccgcagcttcttgatcatctag |

| | | | | |
|---|---|---|---|---|
| pBridge-PIF7mTAD | *PIF7mTAD* | pBridge | cacctcggctgcggcggctgcgactca aagtctcaacggtt | gactttgagtcgcagccgccgcagccgaggtggttggt |
| pBridge-PIF8mTAD | *PIF8mTAD* | pBridge | aggctgcggcggctgcgcatcatcgctc ctcca | atgcgcagccgccgcagcctcatccgtgg |

**Supplementary Table 3. Primers for qRT-PCR analyses**.

| Accession | Gene name | Forward primer | Reverse primer |
|---|---|---|---|
| AT1G69960 | *PP2A* | TATCGGATGACGATTCTTCGTGCAG | GCTTGGTCGACTATCGAATGAGAG |
| AT2G46970 | *PIL1* | AAATTGCTCTCAGCCATTCGTGG | TTCTAAGTTTGAGGCGGACGCAG |
| AT4G16780 | *ATHB2* | TCACAGTACTCTCAATCCGAAGC | CCGTAAGAACTCGCAGTCTAC |
| AT4G14130 | *XTR7* | CACCGTCACTGCTTACTACTTG | CATTGGTGTGAAGAACATAAG |
| AT4G32280 | *IAA29* | CACCATCATTGCCCGTATCA | CCACAGTAGCCGTTGTTGGA |
| AT2G33380 | *RD20* | AAGGACGAAGATGGTTTCCTATC | CGAGAATTGGCCCTCTCTTT |
| AT4G14690 | *ELIP2* | GGCAGAGGCAAAGTCAAAAGG | CGCAACGAGACCGAGCAT |

**Supplementary Table 4. Primer sets for ChIP-qPCR analysis of the *PIL1* locus.**

| Primer set | Forward primer | Reverse primer |
|---|---|---|
| 1 | ATGAATCACGCGGCATTC | ACGTGAGCGGAAAGAACC |
| 2 | GGATGAACAATGCACCACCAC | ACACGAAGGCACCACGAATG |
| 3 | CTCTATGACAGGAACATCACACC | ACAATGACTTGCCTTGTTTACAG |
| 4 | TCGAAGCAAAACCAATCCAAAC | GAATTGTGACCATTCTTTGTTCAG |
| 5 | AGTTGCATTATTGTTGGAGCATC | AAGGTCAGGAAAACACAATGTAG |
| 6 | CTGTGTCATAATCACGATTCAAGG | TTGGGGTTAATGAAGAGCAGC |
